# Toolkits for detailed and high-throughput interrogation of synapses in *C. elegans*

**DOI:** 10.1101/2023.09.12.557402

**Authors:** Maryam Majeed, Haejun Han, Keren Zhang, Wen Xi Cao, Chien-Po Liao, Oliver Hobert, Hang Lu

## Abstract

Visualizing synaptic connectivity has traditionally relied on time-consuming electron microscopy-based imaging approaches. To scale the analysis of synaptic connectivity, fluorescent protein-based techniques have been established, ranging from the labeling of specific pre- or postsynaptic components of chemical or electrical synapses to transsynaptic proximity labeling technology such as GRASP and iBLINC. In this paper, we describe WormPsyQi, a generalizable image analysis pipeline that automatically quantifies synaptically localized fluorescent signals in a high-throughput and robust manner, with reduced human bias. We also present a resource of 30 transgenic strains that label chemical or electrical synapses throughout the nervous system of the nematode *C. elegans*, using CLA-1, RAB-3, GRASP (chemical synapses), or innexin (electrical synapse) reporters. We show that WormPsyQi captures synaptic structures in spite of substantial heterogeneity in neurite morphology, fluorescence signal, and imaging parameters. We use these toolkits to quantify multiple obvious and subtle features of synapses - such as number, size, intensity, and spatial distribution of synapses - in datasets spanning various regions of the nervous system, developmental stages, and sexes. Although the pipeline is described in the context of synapses, it may be utilized for other ‘punctate’ signals, such as fluorescently-tagged neurotransmitter receptors and cell adhesion molecules, as well as proteins in other subcellular contexts. By overcoming constraints on time, sample size, cell morphology, and phenotypic space, this work represents a powerful resource for further analysis of synapse biology in *C. elegans*.

## INTRODUCTION

The nematode *C. elegans* was the first organism for which a full nervous system connectome was established (White et al., 1986). Connectomes now exist for both sexes (Cook et al. 2019; Hall and Russell, 1991; White et al. 1986), the pharynx (Cook et al. 2020; Albertson and Thomson, 1976), for different developmental stages (Brittin et al., 2021; Witvliet et al. 2021), and for the dauer diapause stage (Yim et al. 2023). However, due to the laboriousness of acquiring and analyzing electron microscopy (EM) images and the resulting small sample sizes, our understanding of many aspects of synaptic connectivity has remained limited. For example, we are only beginning to better understand the extent of developmental and inter-individual variability in synaptic connectivity and adjacency (Cook et al., 2023; Brittin et al., 2021; Witvliet et al. 2021) but we remain much in the dark of how exactly external or internal factors establish, alter, and maintain the synaptic connectome. While progress in CRISPR/Cas9 genome engineering and earlier methods continue to increase the array of mutants available to a *C. elegans* geneticist, a lack of synaptic fluorescent reporters and difficulty in scoring them remains a bottleneck for large scale mutant analyses. This is especially true in the synapse-dense regions of the *C. elegans* nervous system, such as the nerve ring.

It has therefore been of paramount interest to establish alternative means to visualize synaptic connectivity that would allow us to score larger sample sizes in living animals. Classically, the study of *C. elegans* synapse biology has been aided by reporters in which various synaptic components are fluorescently labeled (**Figure 1**). For chemical synapses, these include reporters labeling pre- and post-synaptic proteins - such as SNB-1/VAMP, RAB-3, SYD-2/Liprin, CLA-1/Piccolo, GLR-1/AMPAR - fused with a fluorescent protein (Xuan et al., 2017; Mahoney et al., 2006; Yeh et al., 2005; Shen and Bargmann, 2003; Zhen and Jin, 1999; Nonet, 1999; Rongo et al., 1998; Jorgensen et al., 1995) and split-fluorophore transsynaptic technology such as GFP Reconstitution Across Synaptic Partners (GRASP) (Feinberg et al., 2008) and other variants (Feng et al., 2020; Feng et al., 2019), and Biotin Labeling of Intercellular Contacts (iBLINC) (Desbois et al., 2015) (**Figure 1A**). In the case of electrical synapses, innexins can be endogenously tagged with fluorescent proteins to visualize homomeric or heteromeric electrical synapses (Hendi et al., 2022; Gordon et al., 2020; Bhattacharya et al., 2019) (**Figure 1A**). These tools have been somewhat constrained by the relative paucity of cell-specific promoters, but recent single cell RNA-sequencing studies (Taylor et al. 2021, Packer et al. 2019) have yielded novel cell-specific promoters for driving fluorescent proteins in neurons which previously lacked sparse-expressing genes. Moreover, the relative ease of labeling endogenous genetic loci with a fluorophore using CRISPR/Cas9 (Eroglu, Yu, and Derry, 2023; Ghanta et al., 2021; Ghanta and Mello, 2020) has led to an increase in reporters with fluorophore-tagged proteins localized at chemical and electrical synapses (Zhou et al., 2021; Bhattacharya et al., 2019; Lipton, Maeder, and Shen, 2018; Tu et al., 2015; Pinan-Lucarré et al., 2014). With these advances over the years, the *C. elegans* community has built a growing, but still very limited, collection of synaptic reporters. These reporters have been instrumental in studying synapse biology in abundant paradigms including, but not restricted to, synaptogenesis (Mizumoto, Jin, and Bessereau, 2023; Kurshan and Shen, 2019; Jin, 2005), sexual dimorphism (Pechuk et al., 2022; Salzberg et al., 2020; Bayer et al., 2020, 2018; Cook et al., 2019; Hart and Hobert, 2018; Weinberg et al., 2018; Oren-Suissa, Bayer, and Hobert, 2016), and experience-dependent plasticity (Chandra et al., 2023; Bhattacharya et al., 2019; Bayer et al., 2018; Hart and Hobert, 2018). A toolkit of novel reporters is presented in this paper as a resource for the *C. elegans* community (**Figure 1B, Supplementary table 1**).

**Figure 1:**
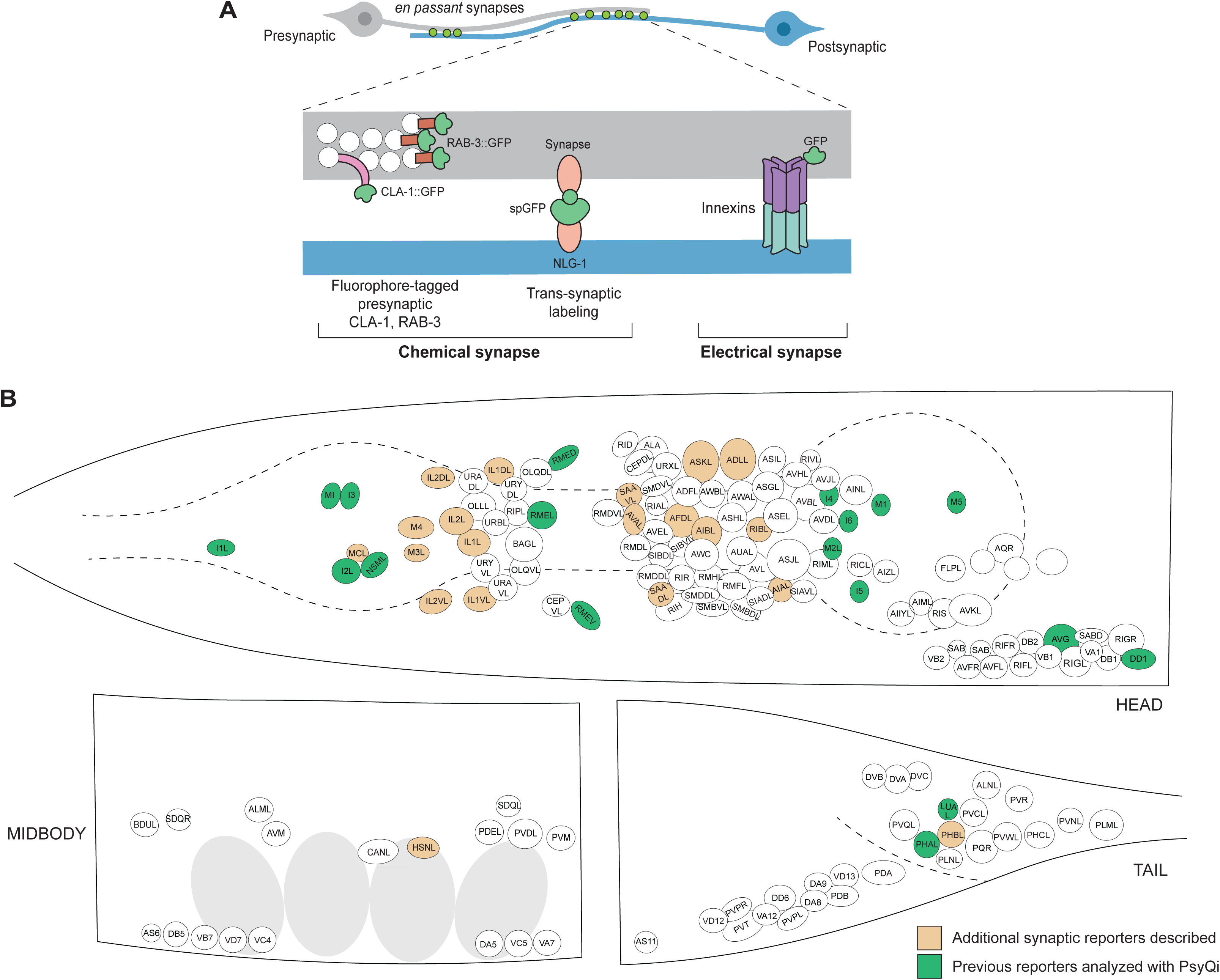
An expanded toolkit of synaptic reporters to study *C. elegans* synapses. **(A)** A schematic of reporters used to visualize *C. elegans* synapses. Chemical synapses were visualized using fluorophore-tagged CLA-1 or RAB-3 proteins to label presynaptic specializations, or synapse-specific reporters using transsynaptic tools such as NLG-1- based GRASP. Electrical synapses were visualized by directly tagging constituent innexin proteins with a fluorophore. **(B)** A schematic showing all neuron classes in the adult *C. elegans* hermaphrodite. Neurons for which new reporters are described in this paper are colored in orange. Neurons for which reporters have been previously published but are quantified with WormPsyQi in this paper are colored in green.

To complement the synaptic reporters and facilitate their proper scoring, we have developed an image analysis pipeline called Worm **<u>P</u>**unctate **<u>Sy</u>**napse **<u>Q</u>**uant**i**fier, or WormPsyQi, which uses machine learning to process and extract useful biological data from a large number of images in an automated, unbiased, and robust manner. By running WormPsyQi on many image datasets of published as well as new synaptic reporters we have developed in the lab (**Figure 1B**), we show that the pipeline can be generalized to synaptic fluorescent reporters of many types, neurons with diverse morphologies, and images taken by multiple users using different microscopes and imaging parameters.

The toolkits described in this paper (strains and imaging pipeline) mitigate several ubiquitous problems related to manual counting of puncta in large datasets. These include, but are not limited to, difficulty in scoring synapses in synapse-dense regions such as the somatic and pharyngeal nerve rings, user bias, human error, problems stemming from scoring images with low signal-to-noise ratio or in regions with high background autofluorescence, laboriousness of scoring large datasets and puncta features such as size and intensity, and inter-dataset variability. We anticipate that the toolkits presented here will provide a motivation for the *C. elegans* research community to continue to build more synaptic reporters and to engage in an analysis of the vast number of presently unstudied synaptic connections in the *C. elegans* nervous system.

## RESULTS AND DISCUSSION

### Generating a semi-automated and robust pipeline for scoring synapses

A key advantage that fluorescent reporters provide over traditional EM-based approaches is that they allow a high-throughput and quick visualization of synapses. However, an ensuing limitation is the quantification of synapse number and ultrastructure. These include plainly obvious features such as the number of synaptic puncta, but also at first indiscernible features such as relative size, fluorescence intensity, and spatial distribution of puncta along a neuron’s processes.

Manual synaptic puncta quantification in large image datasets can be problematic (San-Miguel et al., 2016; Crane et al., 2012). Firstly, it is tedious and low in throughput; for neurons with high synapse density and complex cell morphology in three-dimensional (3D) space, it can take up to several hours to quantify a typical dataset. Secondly, manual quantification requires some a priori information about the region of interest (ROI), thus discouraging observation outside the ROI, which may be of relevance in different genetic or environmental backgrounds. In addition, subtle features may escape manual quantification altogether, possibly hindering their use as a proxy for measuring neuron-specific synapse properties and impeding a comprehensive understanding of pathways which mediate precise synapse distribution. Lastly, manual quantification introduces human error and bias in cases where the fluorescent signal is dim, synapse density is high, the signal-to-noise ratio is poor, or if neurons have complex morphologies which obstruct manual counting in a 3D image stack. These problems are further compounded by dozens of labs generating various synaptic reporters and using distinct microscopes and imaging parameters, all that currently prevents datasets from being easily interpreted, reproduced, shared, and reused. These reasons also partially explain why we cannot rely exclusively on existing image analysis software optimized for other types of datasets or other model organisms.

To address some of these challenges, we have developed a semi-automated computational pipeline, WormPsyQi, to score synaptic puncta in *C. elegans* in a high-throughput manner and with reduced bias. The pipeline is accompanied by a Python-based graphical user interface (GUI) that allows users to seamlessly move through each step and process large volumes of data (**Figure 2 – figure supplement 1B**), which greatly reduces the barrier to use. WormPsyQi consists of five key steps: neurite mask segmentation (optional), synapse segmentation, validation and correction, re- labeling and training (optional), and quantification (**Figure 2A – figure supplement 1A**). First, we obtain a neurite mask. Many image datasets include channels for both synaptic and cytoplasmic markers, and the latter helps exclude noise or autofluorescence signals from other tissues (e.g., the intestine). For neurite segmentation, we modified a convolutional neural network architecture, U-Net (Guo et al., 2022; Guo et al., 2020; Ronneberger, Fischer, and Brox, 2015), to robustly segment the neurite and soma, and create a mask which narrows down the ROI in which synapses are identified in later steps (**Figure 2 – figure supplement 2A**). Masking prior to synapse segmentation is critical if the signal-to-noise ratio is poor and to significantly reduce the overall image processing time by constraining synapse segmentation to the neurite mask.

After masking, a local feature-based pixel classification model segments synaptic pixels inside the neurite mask (**Figure 2 – figure supplement 2B**). In the absence of a neurite mask, the pipeline will indiscriminately segment all puncta within the image. We provide both pre-trained (or default) models and an option for users to train custom synapse segmentation models. Image datasets from four diverse synaptic reporters were used to train four independent default models optimized to detect 1) small synapses (1-3 pixels radius) with low SNR (e.g., puncta with similar intensity and morphology as autofluorescence noise in surrounding tissue), 2) small synapses with high SNR, 3) medium-sized synapses (3-10 pixels radius), and 4) diffuse (>10 pixels radius) signal (see Methods for more detail). These pre-trained models can reliably cover various types of reporters, cells, and imaging parameters as discussed in later sections (**Figures 3-7**). To facilitate the model choice, there is an option for users to test all models on representative images, validate which model works best, and proceed with processing the entire dataset using the optimal model. In addition to this, the GUI enables users to review and edit the synapse segmentation results (**Figure 2 – figure supplement 1C**), as well as to train a custom model by using training data (coupled images and manual annotations of these images – for details, see Github documentation). In our experience, as few as one fully labeled image stack in the training set is sufficient for the segmentation accuracy to converge (**Figure 2C**).

**Figure 2:**
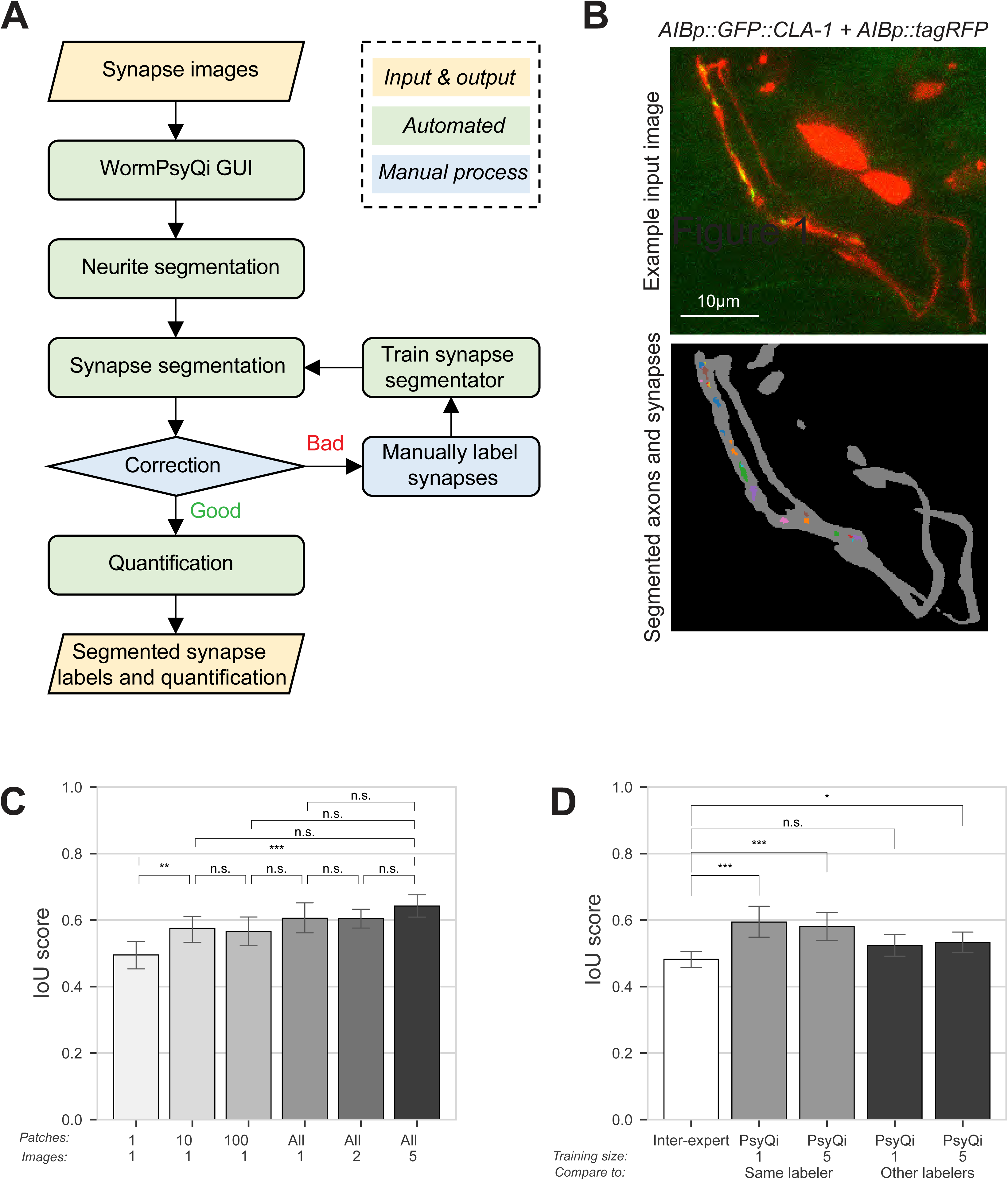
WormPsyQi robustly segments neurites and quantifies synapses. **(A)** A flowchart of the WormPsyQi pipeline. The pipeline is automated except for the correction process, which can be bypassed if the initial segmentation result is good enough. **(B)** Representative input raw and output segmentation images for the interneuron pair AIB. AIB-specific presynaptic specializations and processes were visualized using GFP-tagged CLA-1 (*otIs886*) and a cytoplasmic marker (*otEx8023*). Predicted puncta in the lower panel are colored arbitrarily to represent discrete fluorescent signal. Cell bodies are excluded from the mask to exclude any colocalizing puncta in downstream steps. Images are maximum intensity projections of confocal Z- stacks. **(C, D)** Synapse segmentation accuracy is assessed with IoU score between the ground-truth label and the prediction label. Calculations were performed on six images of the GFP::CLA-1 reporter for neurons I5 (*otEx7503*). The images were labeled by four different experts. **(C)** The IoU score over the size of the training set shows that having one labeled image as a training set is enough to reliably predict synapses for the same conditions of images. In bars 1-3 (from left to right), one image was selected as the training set and then the given number patches in the column label were randomly sampled after the regular patch sampling step of training (see Methods) to further reduce the size of the training set. In bars 4-6, the number of images in the column label were selected as the training set. For each expert’s labels, all possible combinations of the training set were tested and the IoU score for the held-out test set is shown. **(D)** The average IoU score between different experts’ labels is taken as a benchmark (bar 1, leftmost). The IoU scores of WormPsyQi prediction are significantly greater than the benchmark IoU when the same expert’s labels used in the training step were taken as the ground-truth to compare with, both when 1 image and 5 images were used for training. If another expert’s labels were taken as ground-truth, then the IoU score of WormPsyQi prediction was lower but still demonstrated significant improvement compared to the benchmark IoU. *P* values were calculated using one-way ANOVA with Bonferroni correction for multiple comparisons \*\*\**P* ≤ 0.001, \*\**P* ≤ 0.01, \**P* ≤ 0.05 and *P* > 0.05 not significant (ns).

**Figure 3:**
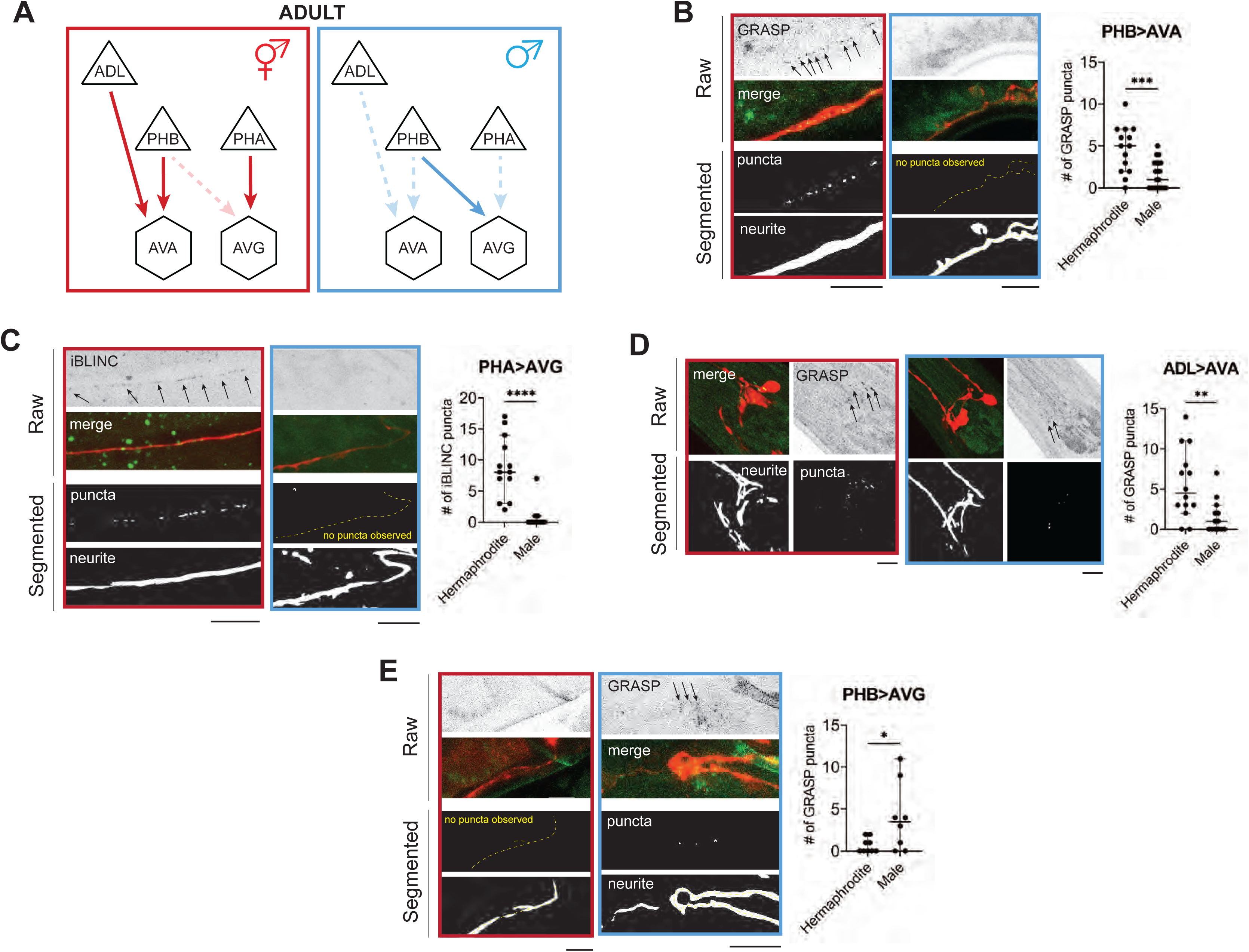
WormPsyQi validates sexually dimorphic synapses in *C. elegans.* **(A)** Subsets of hermaphrodite (red) and male (blue) connectivity diagrams, based on EM studies (Cook et al., 2019; White et al., 1986), showing adult stage sexually- dimorphic synapses analyzed for validating WormPsyQi. Synapses, depicted by arrows, were visualized using GRASP or iBLINC reporters generated either for previous studies (PHA>AVG, ADL>AVA, and PHB>AVG in Cook et al., 2019; Bayer et al., 2018; Oren- Suissa, Bayer, and Hobert, 2016) or this paper (PHB>AVA). Sensory, inter- and motor neurons are depicted as triangles, hexagons, and circles, respectively. **(B)** WormPsyQi validates sex-specific synapses in adult hermaphrodites and males. PHB>AVA, PHA>AVG, ADL>AVA, and PHB>AVG synapses were visualized using the transgenes *otIs839*, *otIs630*, *otEx6829*, *and otIs614*, respectively. Panels corresponding to each reporter show raw confocal images (top) and segmented neurites and synapses (bottom). Segmentation and quantification were performed using WormPsyQi. Synchronized day 1 adult animals were scored. Red – hermaphrodite, blue – male. *P* values were calculated using an unpaired t-test. \*\*\*\**P* ≤ 0.0001, \*\*\**P* ≤ 0.001, \*\**P* ≤ 0.01, and \**P* ≤ 0.05. In each dataset, a dot represents a single worm and lines represent median with 95% confidence interval. All raw and segmented images are maximum intensity projections of confocal Z-stacks. Scale bars = 10μm.

Finally, WormPsyQi also quantifies features of predicted synapses and returns individual and summarized image- and synapse-specific features in a CSV file format, facilitating further statistical analysis. This quantification process not only yields synapse counts (**Figure 3-10**), but also provides metrics more challenging to discern visually. These metrics include total and average synapse volumes, spatial density of synapses, and fluorescence intensity (**Figure 10**). Such comprehensive analysis can enable in-depth analysis across different developmental stages, genetic strains, or imaging conditions. Altogether, WormPsyQi is an ‘image-in-data-out’ pipeline for quantitative analysis of synaptic puncta in fluorescent images.

### WormPsyQi reliably scores synapse features and expedites analysis relative to manual scoring

To evaluate WormPsyQi’s segmentation accuracy and precision, four independent experimenters (**Figure 2D**, inter-expert), manually annotated a dataset of six images, pixel-by-pixel, in which presynaptic specializations of the pharyngeal neuron I5 were labeled with GFP::CLA-1. Training models were constructed from subsets of the dataset and their performance was tested on the remaining images (**Figure 2D**). We selected the intersection over union (IoU) between expert labels and WormPsyQi’s prediction compared to inter-expert label similarity to benchmark the segmentation performance. Notably, the IoU between WormPsyQi’s segmentation and the original labeler’s annotation was significantly higher than the inter-expert IoU (**Figure 2D**, columns 2 and 3), indicating that WormPsyQi allows standardized quantification of synaptic features which human annotators cannot. Importantly, our results show that the WormPsyQi model trained on one image is as good as the one trained on five, indicating that even without using a pretrained model, WormPsyQi can save time spent on annotating synaptic images. When comparing the predictions of WormPsyQi (trained with one expert’s labeling) to the labels of other experts, the IoU scores were found to be comparable to the inter-expert IoU (**Figure 2D**, columns 4 and 5).

To assess the WormPsyQi vs. human labeler effect size, we next sought to compare the discrepancy between WormPsyQi scoring and scoring by two independent human annotators (**Figure 2 – figure supplement 3**). Based on our analysis of I5p::GFP::CLA-1 and M4p::GFP::RAB-3, we found that the effect size was reporter-dependent: for I5 CLA-1, where the puncta were discrete and S/N ratio high, inter-labeler discrepancy and WormPsyQi vs. human (labeler 1 or 2) discrepancy was low. Conversely, the discrepancy across all three scoring methods was high for M4p::GFP::RAB-3, which is a synapse-dense strain with less discrete signal; in this case, semi-automated scoring averaged out human-to-human scoring differences, but did not significantly improve the segmentation.

Lastly, we compared the quantification of additional puncta features such as puncta volume, distribution along the neurite, and overall processing time. To do so, we focused on the ASK GFP::CLA-1 reporter (**Figure 2 – figure supplement 4**). We found that the difference between quantification of puncta number, volume and distribution using WormPsyQi vs. two independent human annotators is statistically insignificant, and that our pipeline reduces quantification time to a large extent (**Figure 2 – figure supplement 4E)**. This is especially true for quantifying subtle features such as size, volume, and intensity, which may not be immediately obvious features and are much more laborious to quantify manually compared to puncta number, because pixel-wise annotation is necessary. Our analysis, therefore, suggests that the algorithm is competent in recapitulating human labeling (in some cases averaging inter-labeler variability) of not just puncta number but also other features, significantly expedites scoring, and can therefore be used to replace manual scoring. The robustness of puncta quantification is further corroborated by comparing WormPsyQi vs. manual quantification for many strains in ensuing sections (**Figure 4 – figure supplement 1, Figure 5**).WormPsyQi vs. manual quantification for many strains in ensuing sections (**Figure 4 – figure supplement 1, Figure 5**).

### WormPsyQi validates sexually dimorphic synaptic connectivity

To demonstrate that our pipeline can quantify synaptic puncta in a robust manner, we initially set out to validate sexual dimorphic synaptic connectivity in the *C. elegans* nervous system. Whole-nervous system EM reconstruction in *C. elegans* adult males and hermaphrodites shows that many ‘sex-shared’ neurons - neurons present in both sexes - display sex-specific, i.e., sexually dimorphic synaptic connectivity patterns (Cook et al. 2019). In previous studies, a small subset of these dimorphic synapses was visualized with GRASP- or iBLINC-based transgenic reporters (Pechuk et al., 2022; Salzberg et al., 2020; Cook et al., 2019; Weinberg and Hobert, 2018; Bayer and Hobert, 2018; Oren-Suissa, Bayer, and Hobert, 2016). By scaling the analyses, presently extremely laborious with EM, these reporters have been indispensable for not only substantiating EM data, but also for understanding the molecular determinants underlying dimorphic circuits and the pathways mediating plasticity (Goodwin and Hobert, 2021 and references therein). In addition, the ability to visualize a specific synapse across many animals continues to reveal new insights such as the extent of variability in synaptic number and spatial distribution across animals, which was previously unappreciated in the absence of tools and sufficient EM samples to study it.

To validate these findings using WormPsyQi, we honed in on four synaptic connections between sex-shared neurons: PHB>AVA, PHA>AVG, PHB>AVG, and ADL>AVA. In all cases, previous studies have shown sex-specific differences in these connections after sexual maturation (**Figure 3A**) (Cook et al. 2019; Oren-Suissa, Bayer, and Hobert, 2016). We show here that WormPsyQi reliably validates previous observations: synapses between ADL>AVA, PHB>AVA, and PHA>AVG are sexually dimorphic, such that synapse number in adult males is less compared to hermaphrodites, and PHB>AVG synapses are less in adult hermaphrodites compared to males (**Figure 3B-E**). An added advantage compared to previous manual puncta quantification was that WormPsyQi expedited the quantification by largely automating the process and, importantly, relied on a single pre-trained segmentation classifier to robustly quantify synaptic number across all four reporters. Together, the performance on these test cases encouraged us to further explore the efficacy of our pipeline in a more systematic manner.

### An expanded toolkit to visualize synapses in *C. elegans*

Having established that WormPsyQi works on a handful of synaptic reporters, we sought to further explore its efficacy in facilitating unbiased synapse quantification. The aforementioned sexually dimorphic reporters (**Figure 3**) are all GRASP- or iBLINC- based and target only 5 neuron classes - most making synapses in the *C. elegans* tail - of the total 116 sex-shared classes in hermaphrodites (White et al., 1986). We asked if our pipeline could quantify various types of punctate signals across other reporter types and in other regions of the nervous system, particularly the nerve ring which comprises the bulk of the hermaphroditic *C. elegans* connectome (Brittin et al., 2021; Cook et al., 2019; White et al., 1986). Despite this preponderance, the nerve ring is remarkably under-studied largely due to a lack of tools available to visualize and reliably quantify synapses in the most complex and synapse-dense region of the *C. elegans* nervous system. This is generally true for synapses in the somatic or central nervous system, compared to neuromuscular junctions, which are easier to visualize and for which relatively more reporters are available.

To broaden the existing resource of synaptic reporter strains, we generated novel synaptic marker strains. These include cell-specific CLA-1 reporters for 8 neuron classes (**Figure 4**) and 5 new GRASP reporters (**Figure 5**) which label specific synapses between multiple neuron classes. Considering both new and previously existing reporters, we describe synaptic reporters for almost 30 neuron classes in both the central and pharyngeal (= enteric nervous systems) of *C. elegans* (**Figure 1; Supplementary table 1**). Many of the new reporters described here target neurons which make synapses in the nerve ring, including cell-specific reporters for IL1, ASK, ADL, RIB, AIA, BDU, HSN (first published in Leyva-Díaz and Hobert, 2022), IL2 (first published in Cros and Hobert, 2022) (**Figure 4**). These efforts relied on the presence of cell-specific promoters driving GFP expression drawing from previous studies. The recent availability of single cell RNA sequencing (scRNAseq) data driven by the CeNGEN project (Taylor et al., 2021) opens up opportunities to gain even broader coverage.

**Figure 4:**
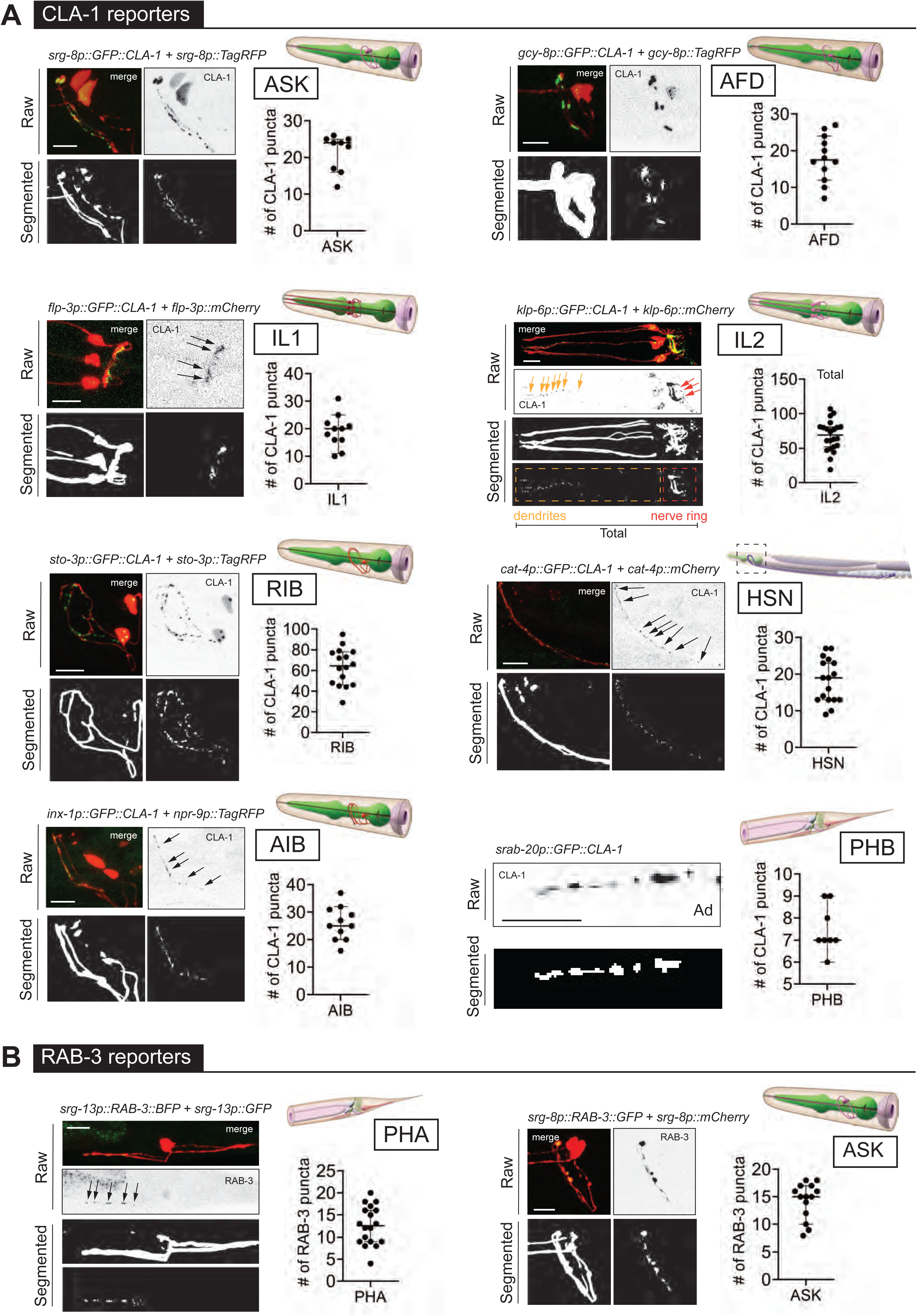
WormPsyQi is generalizable across diverse synaptic reporters targeting the synapses in the central nervous system. **(A)** Representative images of cell-specific synaptic reporters showing presynaptic specializations, as visualized with GFP-tagged CLA-1, for neuron classes ASK (*otIs789*), AFD (*otEx7786*), IL1 (*otEx7363*), IL2 (*otIs815*), RIB (*otIs810*), HSN (*otIs788*), AIB (*otEx8023; otIs886*), and PHB (*otIs883*). **(B)** Representative images of cell-specific synaptic reporters showing presynaptic specializations, as visualized with fluorescently-tagged RAB-3, for neuron classes PHA (*otIs702*) and ASK (*otEx7231*). In each panel, top (raw) and bottom (segmented) images represent both neuronal processes, based on a cytoplasmic marker, and synaptic puncta. There was no cytoplasmic marker in the PHB reporter used, so only puncta are shown. All images are maximum intensity projections of confocal Z-stacks. Quantification of the number of puncta, performed using WormPsyQi, is shown in corresponding graphs. In each graph, a dot represents a single worm and lines represent median with 95% confidence interval. L4 animals were scored unless otherwise noted. Scale bars = 10μm.

**Figure 5:**
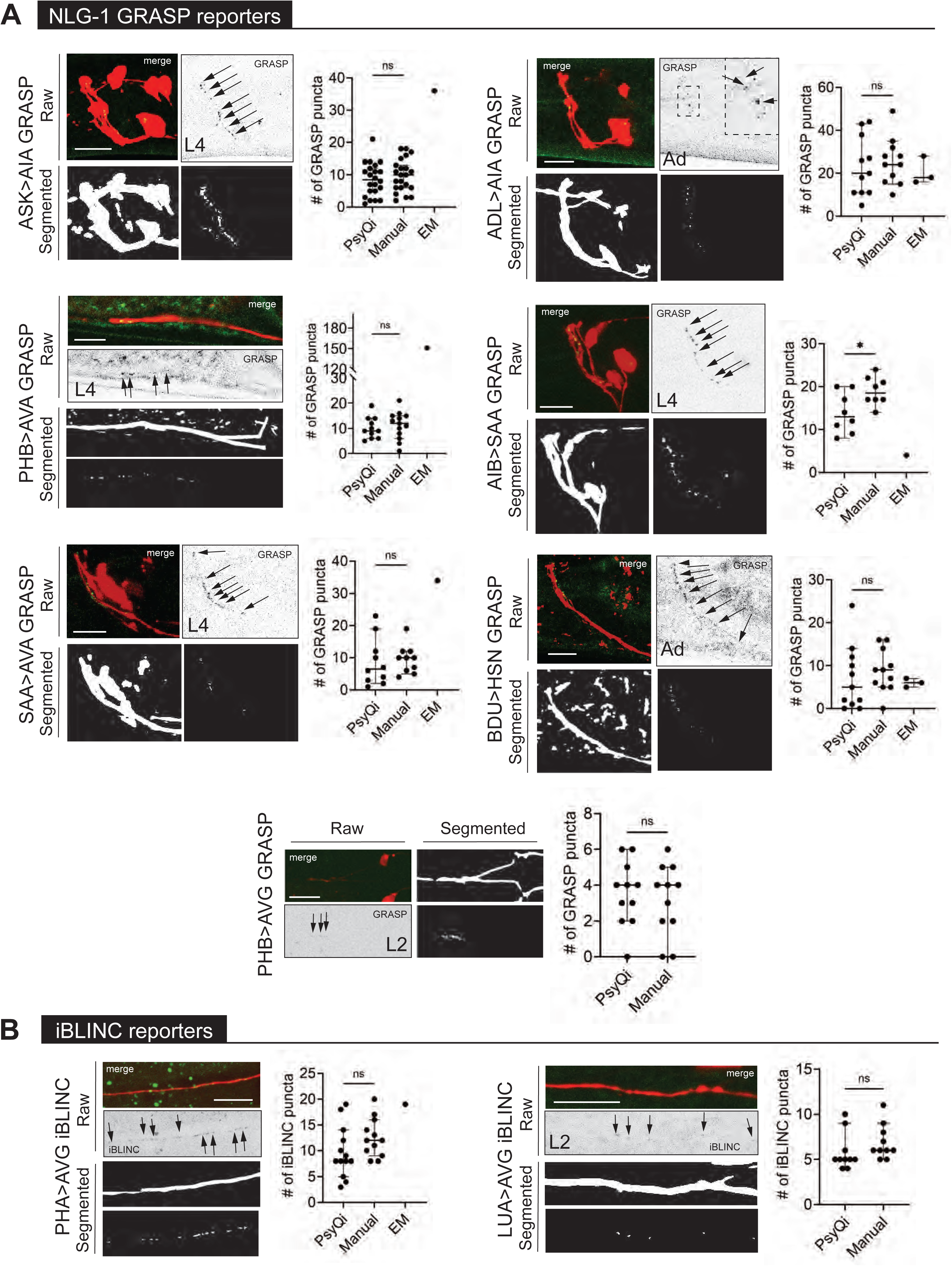
WormPsyQi can quantify synapses in variable GRASP and iBLINC reporters. **(A)** Representative images of NLG-1 GRASP-based synaptic reporters labeling the following synapses: ASK>AIA (*otIs653*), ADL>AIA (*otEx7457*), PHB>AVA (*otIs839)*, AIB-SAA (*otEx7809*), SAA-AVA (*otEx7811*), BDU>HSN (*otEx7759*), and PHB>AVG (*otIs614*). **(B)** Representative images of iBLINC-based synaptic reporters labeling PHA>AVG (*otIs630*) and LUA>AVG (*otEx6344*) synapses. WormPsyQi vs. manual quantification is shown in corresponding plots; EM data is shown where available (Witvliet et al., 2021; Cook et al., 2019; White et al., 1986), but was excluded from statistical analyses due to small sample sizes (n = 1 - 3). In each dataset, a dot represents a single worm and lines represent median with 95% confidence interval. All raw and segmented images are maximum intensity projections of confocal Z-stacks. *P* values were calculated using an unpaired t-test. \**P* ≤ 0.05 and *P* > 0.05 not significant (ns). Scale bars = 10μm.

### WormPsyQi is generalizable across a wide array of synaptic reporters in the central nervous system (CNS)

Previous synapse quantification approaches in *C. elegans* focused on neurons with relatively simple trajectories along the anterior-posterior axis in one plane on the dorso-ventral axis (e.g., *en passant* synapses along the ventral nerve cord). These include semi-automated (San-Miguel et al., 2016; Crane et al., 2012) and manual approaches such as counting synaptic puncta by hand or drawing plot profiles (line scans) of fluorescent signals along a neurite using open-source image processing software such as Fiji/ImageJ, often combined with synapse enrichment quantification as assessed by fluorescence (Kurshan et al., 2018; Xuan et al., 2017; Jang et al., 2016; Dittman and Kaplan, 2006). These methods are well-suited for quantifying synapses along linear neurites but pose problems in neurons with morphologically complex anatomies in 3D space. We thus wanted our pipeline to be able to easily quantify synapses in other types of neurons in *C. elegans*.

In addition, a key observation based on many synaptic reporters imaged by different researchers using diverse microscopes, on different days, and at various life stages and across sexes, is that images collected from these reporters are perceptibly heterogeneous depending on the type of fluorophore-tagged synaptic protein, choice of promoter driving the fluorophore, and the labeling method used. For instance, cell-specific RAB-3 reporters have a more diffuse synaptic signal compared to the punctate signal in CLA-1 reporters for the same neuron, as shown for the neuron pair ASK (**Figure 4 – figure supplement 1C**) and observed previously in other neurons (Cook et al., 2020; Lipton, Maeder, and Shen, 2018; Xuan et al., 2017). This is intuitive based on the distinct localization and function of RAB-3, a presynaptic vesicle protein (Nonet et al., 1997), and CLA-1/Piccolo, an active zone scaffolding protein which recruits synapse vesicles to release sites (Xuan et al., 2017). We note that this is not a consequence of tagging RAB-3 on a specific terminal, since both N- and C-terminal RAB-3 fusions manifest diffuse signal compared to CLA-1. While this may not be a problem for neurons or regions with low synapse density (e.g. ASK RAB-3 and CLA-1 puncta number are similar as shown in Figure 4 – figure supplement 1C), it poses challenges for synapse-dense cases such as AIB and SAA, where WormPsyQi is unable to reliably segment puncta using any of the four pre-trained models. Lastly, within a reporter type, fluorescent signals appear similar, as demonstrated by the similarity of punctate fluorescent signal between various CLA-1 reporters (**Figure 4A**). This is useful because it allows the use of a single pre-trained model for multiple use cases.

In contrast to the relative signal homogeneity in presynaptic fluorophore-tagged reporters of the same kind, signals across reporters that make use of transsynaptic technologies can vary. Fluorescent signal from GRASP and iBLINC reporters appears different from other reporters and is wide-ranging across neurons or synapses (**Figure 5**). As previously reported, this is likely due to transsynaptic labeling methods being more sensitive to the choice of promoters used to drive split-fluorophore fragments, amount of injected reporter DNA, extrachromosomal vs. integrated nature of transgenes, and imaging conditions (Oren-Suissa et al., 2016). Consequently, quantifying the signal of such wide-ranging reporters has been challenging and requires expert knowledge, which is not always transferable across researchers and labs, and introduces human bias and error.

To address these concerns, we used WormPsyQi to quantify puncta in representative datasets for many different presynaptic (**Figure 4**) and transsynaptic (**Figure 5**) reporters for neurons in the central nervous system. We first trained reporter type-specific synapse classifiers (see Methods) to aid quantification so that an average user does not need to perform the tedious step of labeling and training datasets for common types of reporters available in the community. Next, we systematically ran the image datasets through our pipeline, making use of neurite masking (where available) prior to synapse segmentation and quantification (**Figure 2 – figure supplement 1; Figures 4, 5**). Analyzing many reporters further validated that WormPsyQi was able to detect synaptic puncta with minimal human contribution and across a wide range of reporter types (**Figures 4, 5**).

To confirm that WormPsyQi was quantifying fluorescent puncta accurately, we compared its results with manual quantification for a subset of reporters analyzed and found that in most datasets, the difference between WormPsyQi and human annotation was statistically insignificant (**Figure 4 – figure supplement 1A, Figure 5)**. In cases where our pipeline could not recapitulate manual scoring, the masking step was either skipped prior to segmentation (PHB CLA-1) or synapse density was too high (IL1 CLA- 1). In the case of IL1 CLA-1, training a reporter-specific classifier, such as in the case of IL1 (**Figure 4 – figure supplement 1**) improved the results. Generally, having a cytoplasmic reporter in the background of the synaptic reporter enhanced performance; it was also a crucial step in preventing synapse mis-annotation in regions with high background fluorescence intensity such as the tail, where adjacent intestinal autofluorescence typically masks bona fide synaptic signal. This is demonstrated in the case of PHB>AVA GRASP puncta quantification (**Figure 4 – figure supplement 1B**), where WormPsyQi erroneously segments gut autofluorescence puncta as synapses in the absence of a cytoplasmic reporter.

We also found that CLA-1 reporters were overall a superior marker as compared to RAB-3 reporters for the same neuron, as shown for ASK (**Figure 4 - figure supplement 1A, C**). In both cases, WormPsyQi could segment puncta, but the signal was much more discrete, and synapses were resolved better in the ASK CLA-1 reporter. In the case of ASK, overall puncta number in L4 animals was similar in both RAB-3 and CLA-1 reporters but the mean puncta area was much greater in the case of RAB-3, corroborating the diffuse signal which poses quantification challenges for neurons with greater synapse densities, e.g. AIB (**Figure 4 - figure supplement 1D**). We also note that the difference in diffuse vs. discrete signal could not be attributed to the transgenic reporter type (integrant vs. extrachromosomal) since both types of CLA-1 reporters had discrete, smaller puncta compared to RAB-3; owing to variable expressivity in the CLA-1 extrachromosomal reporter, however, fewer puncta could be quantified (**Figure 4 - figure supplement 1C**). Lastly, in some reporters (**Figure 4 - figure supplement 1D**), such as those for RME-specific SNB-1, SAA-specific RAB-3, and AIB-specific RAB-3, WormPsyQi could not effectively segment synapses because the signal was too diffuse and synapse density, in the case of AIB, too high. Again, relying on alternative reporters where possible, such as the CLA-1 based reporter for AIB (**Figure 4A**) which is much better resolved and has distinguishable puncta, is recommended.

In some cases (e.g. AIB>SAA in **Figure 5**), there was discrepancy between WormPsyQi and manual quantification because the signal intensity of puncta was too low to be quantified comprehensively using our pipeline. The analysis of puncta is ultimately influenced by the limits of light microscopy and that the images being analyzed must be of good quality before they are fed into the pipeline. In cases where these issues cannot be resolved during data acquisition, further improvement may be achieved by using alternative microscopy methods, reporter types, or brighter fluorophores. We also note that in several cases, GRASP quantification differed from EM scoring (**Figure 5**). In the absence of a comparable EM sample size in any paradigm tested, it is impossible to assess whether the difference is statistically significant. We note that differences between light-microscopy based imaging and EM are anticipated particularly in synapse-dense regions where our fluorescent reporters, imaging cannot resolve individual synapses as well as EM analysis does.

### WormPsyQi performs robustly in morphologically diverse pharyngeal neurons

In addition to the somatic or central nervous system described thus far, *C. elegans* contains a pharyngeal or enteric nervous system (ENS) which resides entirely in the pharynx, or the feeding organ (Avery and You, 2012; Mango, 2007; Albertson and Thomson, 1976). Unlike the CNS, the dynamics of synaptic connectivity in the ENS are even less explored. Previous EM reconstructions of the pharynx (Cook et al., 2020; Albertson and Thomson, 1976) serve as excellent starting points but, again, we are confronted with limitations such as a very small sample size and a lack of means to study how synapses in the pharynx are established and altered in larval development, as well as under various perturbations.

The *C. elegans* pharynx constitutes 14 neurons classes, and like the ENS in other organisms (Brehmer, 2021; Furness et al., 2007; Costa, Brookes, and Hennig, 2000), there is substantial morphological heterogeneity among these neuron classes (Albertson and Thomson, 1976). In addition, some neurons show variability in branching and synapses, both within a bilateral pair and across individuals, such as in the case of M3 L/R pair (Albertson and Thomson, 1976). This diversity and variability in form poses an additional challenge for image processing, particularly in masking neurites using conventional software. We thus asked if WormPsyQi could create neurite masks of diverse pharyngeal reporters, and subsequently score colocalized synaptic puncta. Cell-specific CLA-1::GFP synaptic reporters of several pharyngeal neuron classes were recently developed, cell-specifically labeling presynaptic CLA-1 in neurons NSM, I1, I2, I5, and MC (Cook et al., 2020). In addition, we describe here two RAB-3::GFP reporters which label presynaptic RAB-3 in neurons M3 and M4 to altogether cover 7 neuron classes in the pharynx (**Figures 6, 9**; **Supplementary table 1**). We also analyzed a pan-pharyngeal reporter in which presynaptic CLA-1 in all 14 pharyngeal classes is visualized with GFP (**Figure 6 – figure supplement 1**) (Vidal et al., 2022).

**Figure 6:**
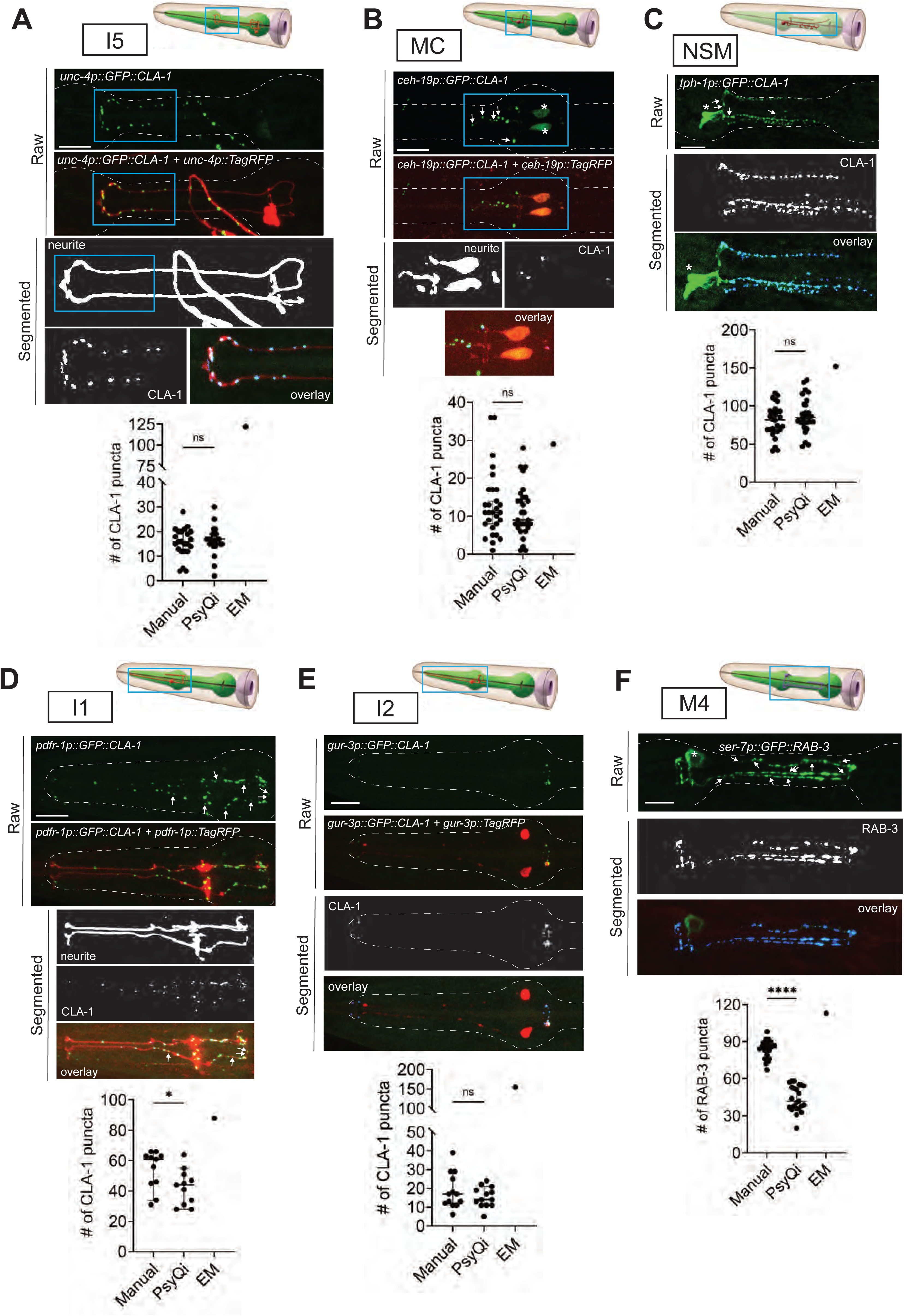
Systematic analysis of pharyngeal synapses using WormPsyQi. CLA-1 or RAB-3 based presynaptic specializations of pharyngeal neurons **(A)** I5, **(B)** MC, **(C)** NSM, **(D)** I1, **(E)** I2, and **(F)** M4 analyzed using transgenes *otEx7503*, *otEx7505*, *otEx7499*, *otEx7501*, and *otIs597*, respectively. For each reporter, a corresponding graph shows manual vs. WormPsyQi quantification, along with EM synapse counts as described in Cook et al., 2020. Day 1 adults were scored, with some datasets previously published (Cook et al., 2020) and re-analyzed here. Masking was used prior to synapse prediction for all neurons except I2, where the cytoplasmic reporter was too dim for creating a continuous mask. Puncta which were not segmented by WormPsyQi are marked with white arrows in the raw GFP image to help inform users what kind of images/reporters are best quantified using WormPsyQi. Cell bodies are marked with an asterisk. *P* values were calculated using an unpaired t-test. \*\*\*\**P* ≤ 0.0001, \**P* ≤ 0.05, and *P* > 0.05 not significant (ns). EM data was excluded from statistical analyses based on limited sample size (n=1). In each dataset, a dot represents a single worm and lines represent median with 95% confidence interval. All raw and segmented images are maximum intensity projections of confocal Z-stacks. Scale bars = 10μm.

Our analyses show that WormPsyQi can robustly segment pharyngeal neurites and generally performs well in quantifying synapses, as with the CNS. This is shown by the comparison between manual and WormPsyQi counts for each reporter (**Figure 6A- F**). In reporters for neurons I5, MC, NSM, and I2, we found no significant difference between puncta count measured by the pipeline versus a human annotator (**Figure 6A- D**). In two reporters, there was a discrepancy between WormPsyQi and human quantification. In the case of the M4 RAB-3 reporter, the RAB-3 signal was too diffuse to resolve all puncta (**Figure 6F**); in reporters for neurons with high synapse densities, such as I1-specific CLA-1 reporter (**Figure 6D**) and pan-pharyngeal CLA-1 reporter (**Figure 6 - figure supplement 1**), some puncta were not detected by the pipeline. In the case of the pan-pharyngeal reporter, WormPsyQi detected most puncta consistently (reflected in varying standard deviation: S.D. = 4.2 for posterior bulb, S.D. = 29.2 for total count) in the synapse-sparse posterior bulb compared to overall puncta for all pharyngeal neurons (**Figure 6 - figure supplement 1**). We expect that presence of a sparse cytoplasmic marker in the background – which would allow using sparse neurite masking before synapse segmentation in WormPsyQi – or alternative labeling strategies would improve results. Finally, we note that there is a marked discrepancy between puncta count in our reporters compared to EM data (**Figure 6**); while the low sample size of EM (n = 1) prevents us from doing any statistical analyses, reasons for plausible differences are explored in the discussion section.

### WormPsyQi can quantify and scale the study of electrical synapses

Although connectomics has historically been biased towards chemical synapses, expression studies on the localization of key electrical synapse components - innexins in invertebrates and connexins/pannexins in vertebrates - suggest the presence of complex connectivity patterns (Ammer et al., 2022; Bhattacharya et al., 2019; Feigenspan et al., 2004; Lee et al., 2003). This is strengthened by evidence of gap junctions in ultrastructural EM-based analyses in both invertebrates (Cook et al., 2019; Jarrell et al., 2012; White et al., 1986) and vertebrates (Smedowski et al., 2020; Anderson et al., 2011), as well as expression studies showing cell type-specific and tissue-specific spatial patterns of innexin and connexin genes (Bhattacharya et al., 2019). In addition, studies involving perturbations of innexins and connexins have shown diverse contributions of electrical synapses to neuronal function (Hendi et al., 2022; Bhattacharya et al., 2019; Hall, 2017; Song et al., 2016; Meng et al., 2016; Kawano et al., 2011; Bloomfield and Völgyi, 2009; Yeh et al., 2009; Chuang et al., 2007).

While the accepted standard for scoring chemical synapses is EM reconstruction, scoring electrical synapses using EM has proven more challenging. EM analysis relies heavily on unambiguous detection of synapse densities; unlike chemical synapses, electrical synapses are more sensitive to EM sample preparation and affected by discrepancy among annotators. Thus, a complete electrical synapse connectome is lacking in any model system to date and remains a difficult endeavor despite ongoing troubleshooting efforts (Emmons, Yemini, and Zimmer, 2021). This makes it even more pressing to develop non-EM based approaches to study gap junctions. Unlike chemical synapses, which are visualized by using transsynaptic labeling or individually tagging pre- and post-synaptic proteins, electrical synapses can be directly visualized by endogenously tagging junction-forming innexins, which assemble into hemichannels in pre- and post-synaptic regions before forming gap junctions via complementary matching between opposing hemichannels (**Figure 1**) (Goodenough and Paul, 2009).

To analyze datasets for electrical synapses using WormPsyQi, we focused on a sparsely expressed innexin reporter, INX-6::GFP (Bhattacharya et al., 2019) (**Figure 7**). WormPsyQi was able to segment and quantify puncta using the default classifier (based on CLA-1 trained images). INX-6::GFP puncta were observed in the pharynx and the nerve ring region, with more puncta scored in dauers compared to well-fed L4 animals b(**Figure 7**). A subset of these puncta represents *de novo* gap junctions between AIB and BAG neurons which mediate dauer-specific chemosensory behavior (Bhattacharya et al., 2019).

**Figure 7:**
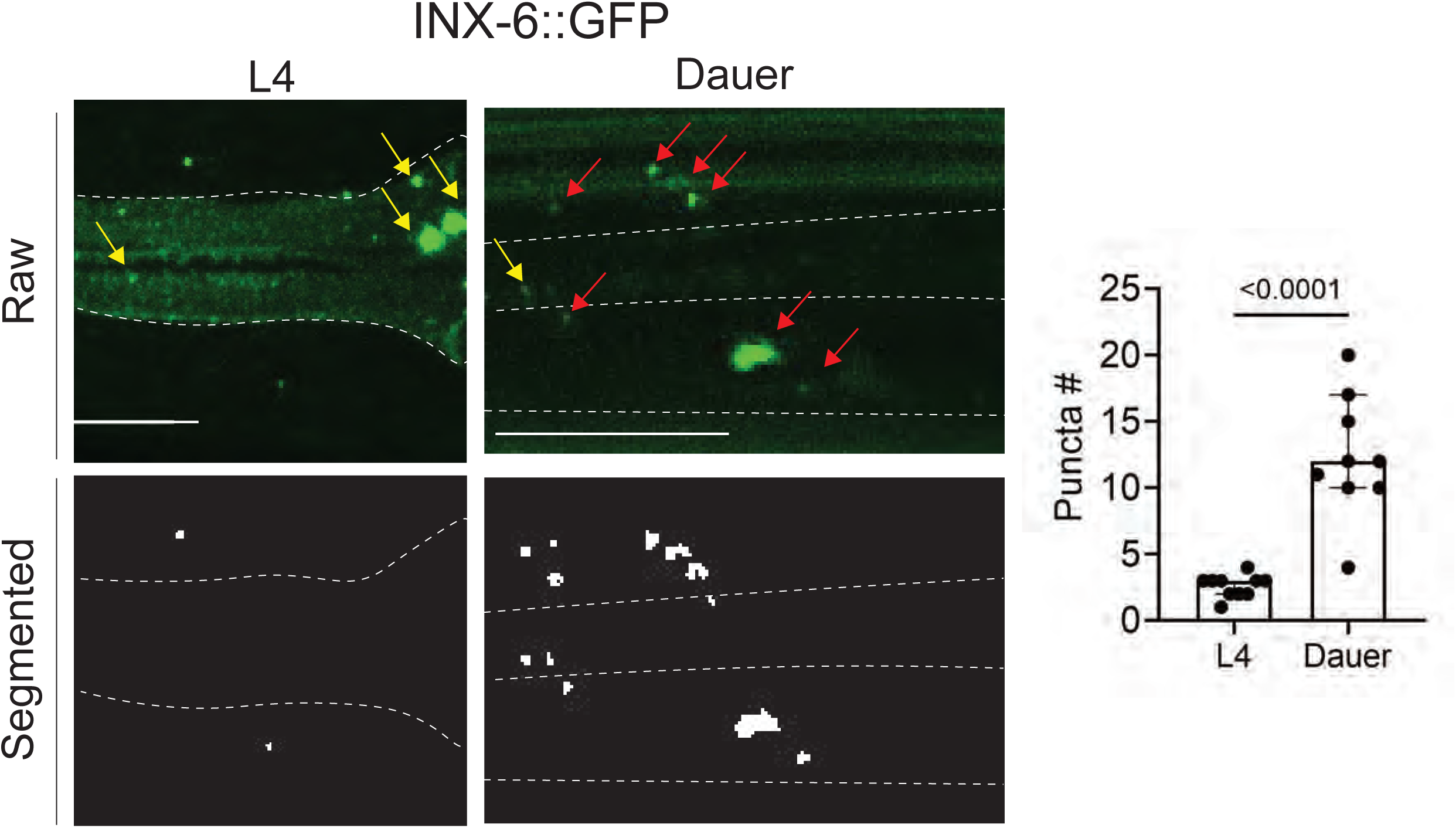
An analysis of electrical synapses using WormPsyQi. INX-6::GFP (*ot805*) puncta are observed in the pharynx and the nerve ring. Nerve ring gap junctions are compared between L4 and dauer animals to validate previous observations (Bhattacharya et al., 2019). Given that outside of the pharynx, *inx-6* is exclusively expressed in AIB interneurons in dauers, the puncta can be assigned to AIB axons as shown previously (Bhattacharya et al., 2019); in the absence of a cytoplasmic marker, WormPsyQi scores a limited number of puncta in L4 animals, but the dauer count significantly exceeds L4 count, as expected. Red arrows denote AIB-localizing electrical synapses and yellow arrows denote electrical synapses in the pharyngeal muscles. All raw and processed images are maximum intensity projections of confocal z-stacks. Statistical analysis was performed using an unpaired t-test. Nerve ring synapses are denoted by a yellow box; synapses outside the box are pharyngeal. In each dataset, a dot represents a single worm and the height of the bar represents median with 95% confidence interval. Images are maximum intensity projections of confocal Z-stacks. Scale bars = 10μm.

### Quantitative analysis of synapses across development in the central nervous system

Synapse plasticity takes multiple forms - such as synapse addition at pre-existing synaptic sites, pruning, and establishment of de novo synapses – and is essential for shaping neuronal circuits and behavior. Intuitively, synapse addition likely prevails circuits building up to meet the needs of a growing animal but this surmise is only recently being confirmed and still remains sparsely understood. Most studies describing synapse addition use EM-based approaches and rely on a small sample size. An obvious pitfall of this is that resulting low-throughput datasets preclude distinguishing synapse plasticity across life stages from baseline inter-individual variability. While expanding EM analysis to study larger sample sizes would be an ideal solution, simply relying on transgenic reporters and the high-throughput analyses they enable could facilitate studying synaptic plasticity in light of variability.

In this paper, we used our toolkits to explore how synaptic puncta change in a small subset of neurons by comparing the number of puncta between early larval and adult stages (**Figure 8**). From our analysis, we found that generally neurons in the CNS progressively add synapses at pre-existing sites throughout larval development up until adulthood (**Figure 8B-E**). For synapses studied in detail (ASK>AIA, PHB>AVA, and AIB>SAA visualized with GRASP, and ASK CLA-1), puncta number steadily increases between early larval and L4 or adult stages. In the case of ASK, synapse addition – shown for all presynaptic specializations visualized with GFP::CLA-1 (**Figure 8B**) and for ASK>AIA synapses (**Figure 8C**), AIA being ASK’s major postsynaptic partner (White et al., 1986) – occurs roughly in scale with the increase in neurite length (**Figure 8A**) in the nerve ring; specifically, there is a ∼2-fold increase across all three measures. In a previous study, a comparative analysis of the nerve ring from samples spanning multiple larval- and adult-staged *C. elegans* showed that systemwide neurite length increases with the length of the animal and that synapses are added progressively thereby maintaining overall synapse density (Witvliet et al., 2021). Since this was done on a systemic scale, albeit with a small sample size, the study noted that synapse addition was likely a cell-specific feature with some neurons adding or pruning more synapses than others (Witvliet et al., 2021). Notably, ‘hub’ neurons added disproportionately more synapses across development. Similarly, a subset of neuron classes that undergo stage-specific plasticity (Witvliet et al., 2021) and/or synaptic pruning with the onset of sexual maturation (some examples are shown in **Figure 3**) remove synapses in a stage- and cell-specific manner.

**Figure 8:**
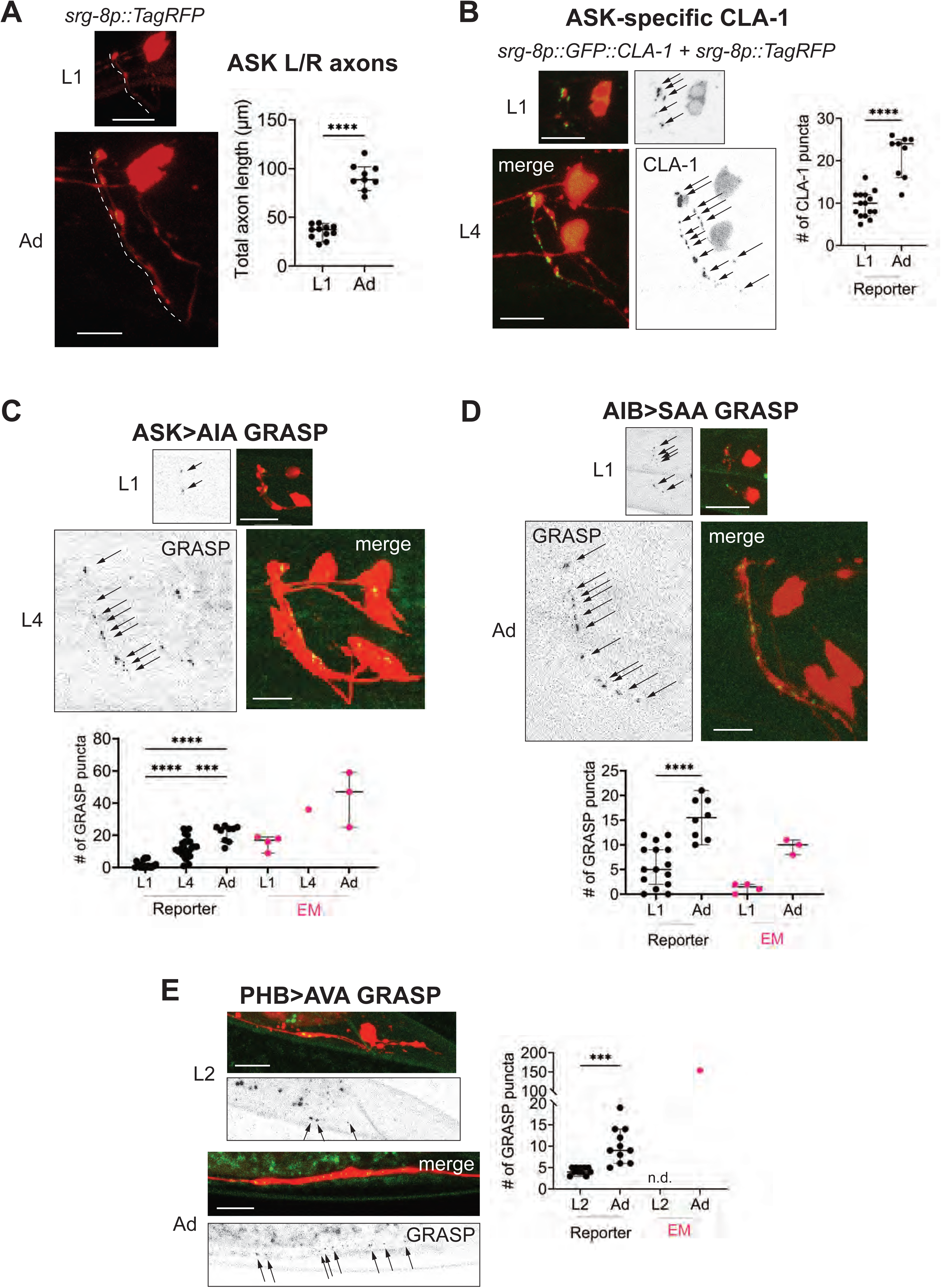
Developmental synapse addition in the CNS roughly scales with neurite length in strains analyzed. **(A)** Representative quantification of the total nerve ring length of late-L1 and day 1 adult animals, as analyzed with an ASK-specific cytoplasmic transgene (*srg-8p::tagRFP* in reporter *otIs789*). Total ASK axon length in the nerve ring increases ∼2-fold between late-L1 and adulthood. Dashed lines mark axons in laterally-positioned worms. **(B)** Representative images of an ASK-specific CLA-1 reporter (analyzed using the transgene *otIs789*) at late-L1 and day 1 adult stages. The graph shows the number of puncta representing ASK-specific presynaptic specializations quantified using WormPsyQi. Number of puncta increases ∼2-fold, which is in proportion with the increase in axon length across development. **(C-E)** Representative images and WormPsyQi-based quantification of synapse addition using synapse-specific GRASP reporters visualizing (**C**) ASK>AIA (*otIs653*), (**D**) AIB>SAA (*otEx7809*) (**E**), and PHB>AVA (*otIs839*) synapses at early larval (L1 or L2) and L4 or adult (Ad) stages. *P* values were calculated using an unpaired t-test **(A, B, D, E)** or one-way ANOVA with Bonferroni correction for multiple comparisons **(C)**. \*\*\*\**P* ≤ 0.0001 and \*\*\**P* ≤ 0.001. EM data (taken from Witvliet et al., 2021; Cook et al., 2019; White et al., 1986) was excluded from statistical analysis based on limited sample sizes (n = 1 - 4) which were out of proportion with reporter-based data. In each dataset, a dot represents a single worm and lines represent median with 95% confidence interval. All raw and segmented images are maximum intensity projections of confocal Z-stacks. Scale bars = 10μm.

Uniform synapse addition has also been noted in sparse EM-based comparative connectomic studies in other systems, such as the nociceptive circuits in *D. melanogaster* between first and third instar larval stages (Gerhard et al., 2017), motoneurons in *D. melanogaster* across larval development (Couton et al., 2015), antennal lobe in *D. melanogaster* between late pupa and adulthood (Devaud et al., 2003), and primary somatosensory and visual cortical regions in mice and rhesus macaque during early postnatal development (Wildenberg, Li, and Kasthuri, 2023). Importantly, since our approach is high-throughput, it allows us to ascertain synapse addition while accounting for inter-individual variability in synapse number in the reporters studied. This is a key point which remains hard to address with current EM output, but focusing on a ‘core’ set of synapses present across EM samples (Cook et al., 2023) presents an opportunity for statistically-powered analyses.

### Quantitative analysis of synapses across development in the enteric nervous system

Synapse addition may be widespread but it is not ubiquitous when studied at the level of neuron class or region of the nervous system. To broaden our analysis of developmental plasticity in *C. elegans*, we next focused on neurons of the ENS. The length of the pharynx from the tip of the corpus to the end of the terminal bulb increases 2-fold (from 60μm to 130μm, roughly) between L1 and adulthood (Shibata et al., 2016); this elongation is modest relative to the overall ∼4.5-fold increase (from 250μm to 1150μm, roughly) in the length of the animal across the same developmental window (Witvliet et al., 2021; Shibata et al., 2016) (**Figure 9A**). To investigate how neuronal processes and presynaptic specializations of pharyngeal neurons scale with the elongation of the whole organ, we quantified neurite length (denoted by ‘Length of ROI’) and the number of fluorescently-tagged CLA-1 or RAB-3 puncta in the pharyngeal neuron classes MC, M3, I2, M4, and NSM (**Figure 9B-F**). In this case, puncta were counted both manually (**Figure 9**) and using WormPsyQi (**Figure 9 – figure supplement 1**). Due to high synapse density and lack of or faint expression of a cytoplasmic reporter in most strains analyzed, WormPsyQi undercounted puncta in early larval stages (**Figure 9 – figure supplement 1**). However, trends in developmental plasticity between manual and automated quantification were similar (**Figure 9 – figure supplement 1**).

**Figure 9:**
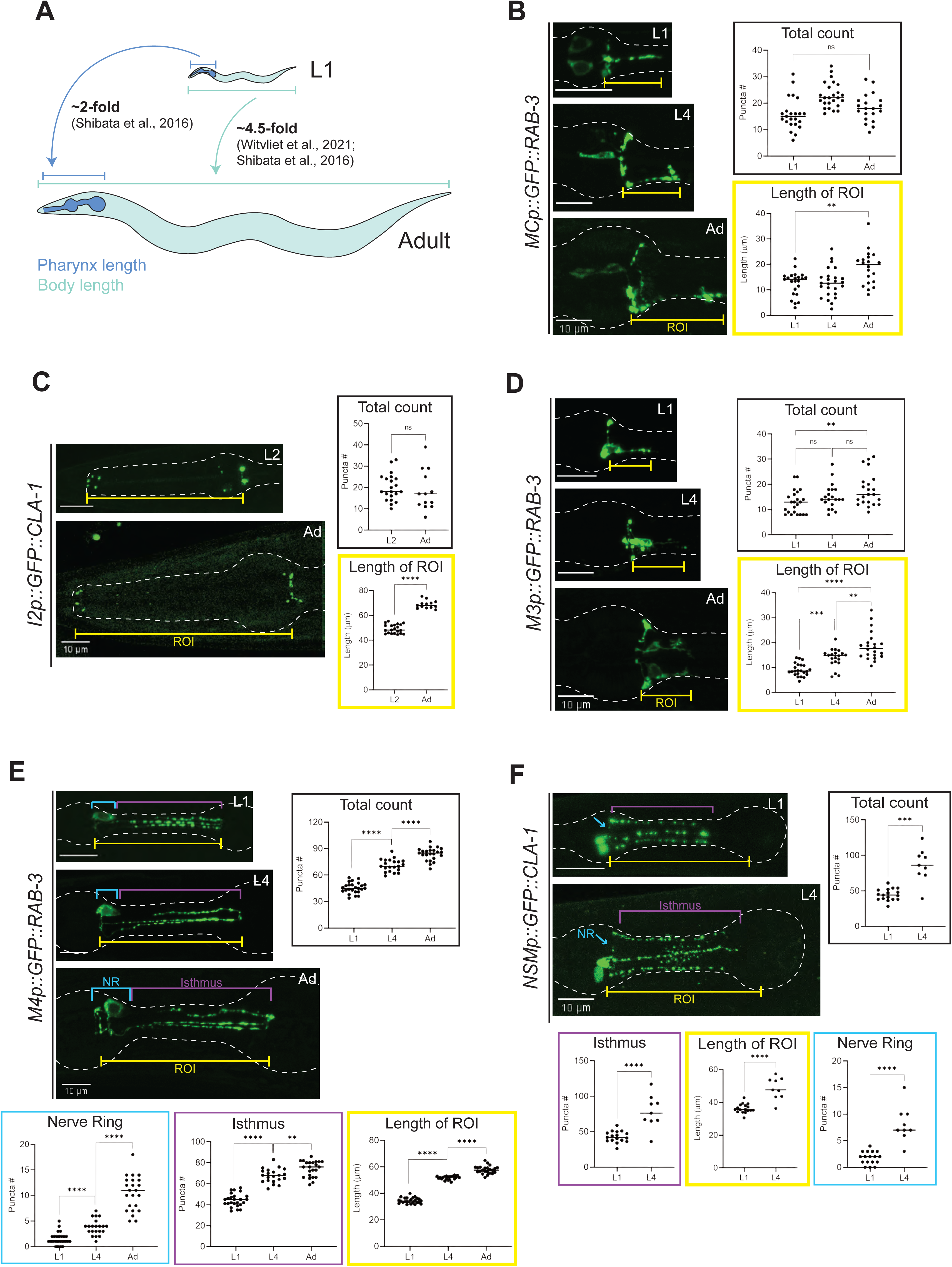
Synapse addition in the enteric nervous system does not scale with neuron growth over developmental. **(A)** Overview of pharynx and total body length elongation between L1 and adult stages based on previous findings (Witvliet et al., 2021; Shibata et al., 2016). **(B-F)** Representative images and quantification of number of presynaptic specializations at early larval and adult stages visualized using **(B)** GFP::RAB-3 in MC (*otEx7505*), **(C)** GFP::CLA-1 in I2 (*otEx7497)*, **(D)** GFP::RAB-3 in M3 (*otIs602*), **(E)** GFP::RAB-3 in M4 (*otIs597)*, and **(F)** GFP::CLA-1 in NSM (*otEx7499*) neurons. Manual counts of total CLA- 1 or RAB-3 puncta are plotted for each reporter (black boxes). For neurons MC, M3, M4, and NSM, region of interest (ROI, yellow box) length was measured from the pharyngeal nerve ring to the most posterior CLA-1 or RAB-3 signal observed. For I2 neurons, ROI length was measured from the pharyngeal nerve ring to the tip of the anterior end of the pharynx based on brightfield. For NSM neurons, ROI length was measured from the pharyngeal nerve ring to the grinder in the terminal bulb, based on brightfield. For neurons M4 and NSM, additional quantification was performed for the pharyngeal nerve ring region (NR, blue box) and the isthmus (purple). *P* values were calculated using Mann-Whitney test. \*\*\*\**P* ≤ 0.0001, \*\*\**P* ≤ 0.001, \*\**P* ≤ 0.01, and *P* > 0.05 not significant (ns). In each dataset, a dot represents a single worm and lines represent median. All images are maximum intensity projections of confocal Z-stacks. Scale bars = 10μm.

We found that in contrast to neurons in the CNS, neurons in the ENS show striking disparity in synapse addition across neuron classes. The number of CLA- 1/RAB-3 puncta in neurons MC, I2, and M3 remains constant (neurons MC and I2, **Figure 9B, C; Figure 9 – figure supplement 1A, D**) or increases modestly compared to the elongation of the pharynx (neuron M3, **Figure 9D**; **Figure 9 – figure supplement 1B**) between early larval stages (L1 or L2) and adulthood. This is in spite of the neurite length scaling roughly in line with the length of the pharynx in all three cases (**Figure 9B-D**). In other words, synapse density in these neurons decreases across development. We note that in some neurons, such as M3 (**Figure 9D**) (Albertson and Thomson, 1976), it is possible that the large variability in synapse number across individuals and within a bilateral pair, at a given stage in development, may complicate comparative analysis between stages. Furthermore, since our analysis does not include quantification of neuronal adjacency across development, we cannot rule out the possibility that physical contact within neighborhoods of pharyngeal neurons which make synapses with one another is not scaling with the elongation of the pharynx or the whole animal. EM reconstructions, because of their often system-wide nature, may be better-suited for this type of analysis since it yields quantification of both the connectome and the adjacencies, or ‘contactome’, between synaptic partners.

In contrast to neurons which show variable synapse density across ages, we found two neuron classes, M4 and NSM, where CLA-1/RAB-3 puncta are added proportionately with increasing neurite length and overall pharynx elongation across development (‘Total count’ in **Figure 9E, F; Figure 9 – figure supplement 1C**). Our CLA-1-based data for NSM also corroborate previous findings of the developmental trajectory of NSM’s synaptic branches (Axäng et al., 2008). Based on previous EM studies (Cook et al., 2020; Albertson and Thomson, 1976), these puncta largely correspond to neuromuscular junctions, many of which arise post-embryonically and can be considered somewhat separate from the ‘core’ pharyngeal connectome established in the embryo which seems more static. To take a deeper look, we calculated M4 and NSM puncta number separately in the pharyngeal nerve ring and isthmus, and we found synapse addition between L1 and adulthood in both regions (**Figure 9E, F**). Given the density of puncta in the nerve ring, it is ultimately difficult to discern change in neuron-to-neuron synapses made by these neurons by relying solely on CLA-1/RAB-3 reporters. Synapse-specific reporters built using GRASP/iBLINC or temporal EM analysis will offer greater resolution needed to fully understand developmental plasticity in these synapse-dense regions. In conclusion, synapse addition appears to differ between the ENS and CNS, as suggested by the lesser increase in synapse number in pharyngeal neurons relative to somatic neurons in our data (**Figures 8, 9**).

### Using WormPsyQi to extract subcellular information of synapse distribution

Next, we exploited PsyQi to extract subcellular information of synapse distribution. To this end, we quantified spatial positions of presynaptic specializations in neurons PHA, AIB, and ASK as proof of principle (see Methods). We found that the presynaptic reporter data recapitulated expected subcellular distribution patterns in all three neurons analyzed. Previous EM data (Cook et al., 2019; White et al., 1986) shows that in PHA neurons, presynaptic sites are concentrated - and uniformly distributed – in the axonal region anterior to the branch point with close to zero synapses posterior to it; our PHA-specific RAB-3 reporter recapitulates this topological distribution (**Figure 10A**). Taking advantage of our sizeable dataset (n = 17 compared to EM, n = 1) which represents an ‘average’ population of worms, we took the analysis a step further and measured the probability of finding a synapse at a particular location along the axon, thus quantitatively showing the distribution of stereotyped (high probability density) and variable or noisy (low probability density) sites; we found that distribution was indeed more or less uniform in the case of PHA in spite of inter-individual variability (**Figure 10A**, **Figure 10 – figure supplement 10A**).

**Figure 10:**
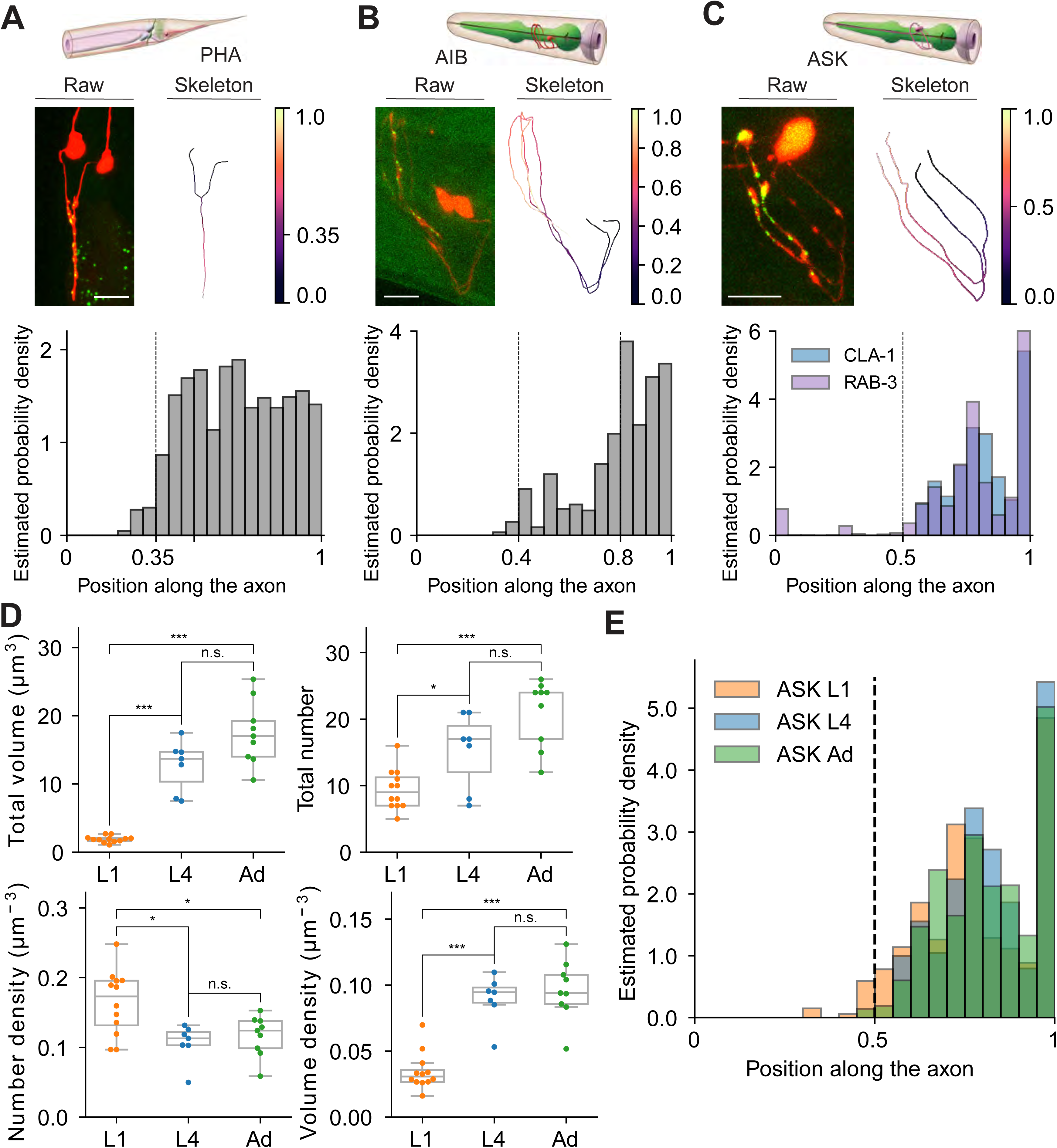
Extracting subcellular and spatial synaptic features using WormPsyQi. (**A-C**) Illustration of distribution profile of RAB-3 puncta in PHA, CLA-1 puncta in AIB, and CLA-1 and RAB-3 puncta in ASK neurons. Top: Schematic diagram of neurons (source: WormAtlas). Middle left: raw image data with contrast enhancement and maximum projection. Middle right: skeleton diagram of neurons. Position 0 indicates SOMA and 1 indicates the furthest axon terminal. Bottom: Probability density histogram of puncta distribution along the neurite process. Distribution profiles of worms at the same stage were combined to give a histogram (see Methods). (**C**) RAB-3 and CLA-1 puncta profiles are overlaid. (**D**) Example features describing developmental changes of ASK neuron’s connections. Top: puncta volume and puncta number per worm, from left to right. Bottom: puncta number density and puncta volume density per worm (calculated along the neurite domain covered by synapses). *P* values were calculated using one-way ANOVA with Bonferroni correction for multiple comparisons. \*\*\**P* ≤ 0.001, \*\**P* ≤ 0.01, \**P* ≤ 0.05 and *P* > 0.05 not significant (ns). In each dataset, a dot represents a single worm. (**E**) Synaptic puncta density distribution in ASK neurons at different stages. For each worm, the synaptic volume distribution is normalized by axon length.

We performed a similar analysis on CLA-1 based reporters for neurons AIB and ASK (**Figure 10**). AIB’s axons occupy two distinct neighborhoods of the nerve ring (Moyle et al., 2021), anterior and posterior, with a majority of presynaptic sites localizing in the anterior region (Sengupta et al., 2021). Both RAB-3 (Sengupta et al., 2021) (**Figure 4 – figure supplement 1D**) and CLA-1 (**Figure 4A**, **Figure 10B**) reporters for AIB qualitatively show this distribution. Quantification of topological synapse distribution within the anterior axon in a representative population revealed that synapse density is sparse in the ventral regions, but increases gradually as the processes travel to the crossover point at the dorsal midline, with highest density proximal to it. We also observed widespread variability in puncta size that could not be correlated with spatial location (**Figure 10 – figure supplement 1B**). Lastly, in the case of ASK neurons, EM data shows (White et al., 1986) distinct regions along the axons where synapses localize. Our RAB-3 (**Figure 4B**) and CLA-1 (**Figure 4A**, **Figure 10C**) recapitulate this distribution pattern in a representative population. We found that ASK axon regions differed markedly in absence or presence of a synapse as well as stereotyped vs. variable sites as denoted by the distinct peaks in the estimated probability density plot (**Figure 10C**).

Having earlier quantified the developmental plasticity of synapse number in ASK neurons between L1 and adulthood (**Figure 8**), we next asked how the stereotyped topological distribution and synapse ultrastructure features (such as synapse volume) change across development. The total volume of synapses scaled proportionately with synapse addition between L1 and adult animals (**Figure 10D**), and peaks in probability density were stereotyped as well (**Figure 10E**), suggesting that these ultrastructure and spatial quantifications followed the same extent of plasticity as the overall number of synapses did. In addition, we plotted the number of puncta against ultrastructure features including mean volume and mean intensity, and found that mean volume, in the ASK>AIA reporter studied in detail, clustered in an age-dependent manner; conversely, the synapse density vs. mean intensity feature space did not show any age-dependent clustering (**Figure 10 - figure supplement 2**).

Although this analysis is limited to a few neurons, the observation that synapse distribution - for synapses that are present at birth - remains stereotyped is not unanticipated given that, aside from inter-individual variability, the overall architecture of the brain in terms of cell body positions, neuronal morphologies, and relative arrangement of processes within neighborhoods in *C. elegans* are largely maintained across development (Witvliet et al., 2021; Moyle et al., 2021; Brittin et al., 2021). An exception to this organizational stereotypy is a subset of neurons that undergo drastic changes across one or several of these spatial features between L1 and adulthood, such as birth of new neurons (e.g. VNC neuron classes), migration of cell bodies (e.g. HSN, ALM), dendritic growth (e.g. FLP, PVD, NSM), synapse pruning (e.g. sexually-dimorphic neurons PHB, AVA etc.), and neighborhood change (e.g. DD and VD neurons) (Androwski et al., 2020; Emmons, 2018; Howell, White, and Hobert, 2015; Middelkoop and Korswagen, 2014; Altun and Hall, 2011; Oren-Suissa et al., 2011; Smith et al., 2010; Axäng et al., 2008). Overall, precise quantification of synapse distribution patterns and ultrastructural features on a population level, something that can only be done qualitatively by eye, is a useful way of assigning topological information to reporter-based datasets and can be valuable readouts under various genetic and environmental perturbations, especially in cases where the synapse number may remain unchanged and are uninformative for phenotyping.

### Limitations of the toolkits described

We have demonstrated that the toolkits described here enable a high throughput, unbiased, in-depth, and robust workflow for studying chemical and electrical synapses in *C. elegans* based on analyzing many reporters. However, we note some important limitations. First, the reporters presented here are a proxy for chemical synapses and may represent simply components of synapses. For instance, in the case of GRASP - which relies on split GFP fragments fused usually with Neuroligin/NLG-1 - when two adjacent neuronal processes come in proximity to form a synapse, the fragments are reconstituted and result in fluorescent puncta. Although these puncta suggest synapses and the tool has been instrumental, it is important to note that GRASP specifically visualizes NLG-1 localization and hence the interpretation of puncta number and features quantified with WormPsyQi categorically represent NLG-1 features and dynamics. Intuitively, GRASP would not be ideal for neurons which do not endogenously express NLG-1 at the right time or localize it to synaptic domains, therefore under- or mis-representing the synapse population between two neurons.

Another limitation of the reporters described is that they contain over-expressed multicopy transgenic arrays which could lead to trafficking and localization artifacts. While some of these problems are mitigated by injecting transgenes at different concentrations, integrating extrachromosomal arrays, and testing multiple gene promoters to drive expression, these measures occasionally fail and may introduce a selection bias. In the case of transsynaptic technologies where multiple transgenes need to be co-injected, artifacts tend to be harder to identify and eliminate. Comparisons of multiple reporters expressing an identical transgene often show variability across lines picked from the same injection. This holds true for both presynaptic and transsynaptic reporters although the latter tend to have more variability across lines. This could partially explain discrepancies between data from transgenic reporters and EM reconstructions (**Figures 5, 6, 8**). With gene editing becoming easier, an increasing collection of reporters with endogenously-tagged proteins known to localize in specific neurons and at specific sites along a neurite will add to our knowledge of pre- and post-synaptically localized proteins available to visualize synapses. Reporters with endogenously-tagged proteins and single-copy transgenes show more consistent expression and spatial localization. Indeed, the use of diverse types of reporters, when available, will be essential to establish the ground truth and inform mutant analyses for advancing our understanding of synapse biology.

Lastly, although we have shown that the image analysis pipeline is robust and versatile, it is constrained by the type of images and reporters being analyzed. In cases where the cytoplasmic marker is suboptimal (e.g. dim cytoplasmic marker in CLA-1 reporters for pharyngeal neurons I2 and NSM, or dim signal on the far end of the cover slip while acquiring images), the resulting neurite mask may be discontinuous, resulting in undercounting of synaptic puncta in downstream steps. In such scenarios, it may be useful to visualize the neurite mask in Fiji prior to synapse quantification. Since this is technical constraint independent of WormPsyQi but can directly affect its performance, we suggest resolving this prior to processing an entire dataset. For instance, one can use a stronger promoter to drive the cytoplasmic fluorophore, perform some preprocessing such as deconvolution, denoising, or contrast enhancement prior to synapse segmentation, or skip the masking step altogether and directly segment synapses. The last scenario is particularly useful in cases where the synapse signal is brighter than the cytoplasmic reporter; for our quantification, we have relied on this for several reporters and denoted them with “no mask” in Supplemental Table 1. We note that the processing time is significantly reduced in the presence of a mask, so if the user decides to skip masking prior to synapse segmentation, we suggest cropping the precise ROI so the quantification can be expedited. Also, where the synapse signal is diffuse (e.g. RAB-3 reporters) or synapse density is too high (e.g. CLA-1/RAB-3 reporters for IL2, NSM, AIB etc., **Figures 4, 6**), quantification can be inaccurate. Analyzing the “overlay” images saved during image processing can reveal problematic regions of interest and help the user decide whether to proceed with quantification. In general, if the fluorescent signal in an image is visibly discrete, WormPsyQi works well in quantifying many discernible and subtle features which would otherwise be tedious or impossible to achieve.

## MATERIAL AND METHODS

### Strains and maintenance

Wild-type strains used were *C. elegans* Bristol (strain N2). Worms were grown at 20°C on nematode growth media (NGM) plates seeded with bacteria (*E.coli* OP50). Details of all transgenic strains are listed in **Supplemental table 1**; corresponding sex and age are also noted.

### Cloning and constructs

All constructs used to build the synaptic reporters are listed in **Supplemental table 2**. In most cases, subcloning by restriction digestion was performed to generate GRASP, CLA-1, and cytoplasmic (TagRFP, mCherry) constructs. Cell-specific promoter fragments were amplified from N2 genomic DNA and cloned into 5’ SphI- and 3’ XmaI-digested vectors expressing fluorophore only (in the case of cytoplasmic constructs), GFP::CLA-1, GFP::RAB-3, and split-GFP (spGFP1-10, spGFP11) sequences; T4 ligation was used to insert digested promoter fragments into digested vectors. A plasmid containing GFP::CLA-1(S) (PK065) was kindly provided by Peri Kurshan. In some cases, RF-cloning or Gibson Assembly (NEBuilder HiFi DNA Assembly Master Mix, Catalog # E2621L) were used as the preferred choice of cloning. Primers for specific promoter fragments are listed in **Supplemental table 2**.

Transgenic strains were generated by microinjecting constructs as simple extrachromosomal arrays into N2 or *him-5 (e1490)* genetic backgrounds. Precise concentrations of co-injected constructs are detailed in **Supplemental table 2.** Extrachromosomal array lines were selected according to standard protocol. Where integration was performed, gamma irradiation was used.

### Microscopy and image analysis

Worms were anesthetized using 100mM sodium azide (NaN3) and mounted on 5% agar on glass slides. Worms were analyzed by Nomarski optics and fluorescence microscopy, using a 40x or 63x objective on a confocal laser-scanning microscope (Zeiss LSM880 and LSM980) or a spinning disk confocal (Nikon W1). When using GFP, we estimated the resolution of our confocal to be ∼250 nm. 3D image Z-stacks were converted to maximum intensity projections using Zeiss Zen Blue or ImageJ software (Schindelin et al., 2012). Manual quantification of puncta was performed by scanning the original full Z-stack for distinct dots in the area where the processes of the two neurons overlap. Figures were prepared using Adobe Illustrator.

### Statistical analysis

Statistical tests were performed using Python (version 3.10) or GraphPad PRISM (version 9.5.1). Where multiple tests were performed, post-hoc Bonferroni correction was used to adjust P values for the number of pairwise comparisons made.

### Neurite segmentation

#### Model structure

We adapted a ‘2.5D’ U-Net model (Guo et al., 2020) to segment neurites from 3D fluorescent image stacks. The input is a 3D chunk with dimension 7 x 256 x 256 (D x H x W), and the prediction of the center slice (256 x 256). The chunk that includes nearby slices provides more spatial context compared to 2D segmentation, while reducing the memory constraints of 3D segmentation. The model has an encoding block and a decoding block with skip connections at the same resolution level that “squeeze” 3D into 2D (**Figure 2 - figure supplement 2A**). The structure of the encoding block is detailed in Figure 2 – figure supplement 2A’.

#### Training process

The U-Net model was trained using a 3D fluorescent microscopy dataset. The dataset covers 5 neurons from different developmental stages (ASK, AIA, PHB, AVA, I5), with 16 images in total. We split the dataset into the ratio 10:2:4 for training, validation, and testing, respectively. Due to the extremely large size of the original image stack, we sampled each 3D stack randomly into 200 smaller 3D patches as stated above and filtered out the blank background patches using a threshold on mean intensity. For the training process, we used a combination of IoU loss (Berman, Triki, and Blaschko, 2018) and the cross entropy loss: L = L_IoU_ + L_CE,_ with the learning rate set to 0.001. The training usually converges within 200 epochs. Data augmentation (including random flipping and rotation), is added for each epoch. The detailed benchmark summary is shown in **Supplemental table 3**. All training experiments were done using the same dataset with 4-fold cross validation, and same loss function and batch size. The metrics shown in the table refer to the neurite class, except for accuracy, which shows an overall pixel prediction accuracy. Training and inference were done on a Nvidia V100 16GB GPU.

### Synapse segmentation

#### Process overview

To segment synapses from 3D fluorescent image stacks, WormPsyQi uses a two-layer pixel classification model (**Figure 2 – figure supplement 2B**). In the first layer, a set of predefined 2D and 3D local filters are applied to each pixel to generate a feature vector. The first-layer classifier then estimates the probability that a pixel belongs to a synapse based on this feature vector. In the second layer, the probability estimations of a pixel and its nearby pixels are added to the first layer’s feature vector to create an extended vector. This extended vector is then fed into the second-layer classifier to determine whether the pixel belongs to a synapse. Optionally, the binary neurite mask from the neuron segmentation step is used to filter out the false-positive pixels that are not in the vicinity of synapse-forming regions. Although we primarily used the Random Forest for the classifiers in this paper, WormPsyQi provides Random Forest (RF), Support Vector Machine (SVM), and multilayer perceptron as options for training a new model with the GUI. While SVM is effective in many different machine learning applications and generally less prone to overfitting with a small dataset, it is sensitive to noise and its fit time complexity is more than quadratic, making it less ideal for large training datasets. During training, grid search hyperparameter optimization with 5-fold cross-validation is used. For prediction, if the target image has a neurite mask segmented in the previous step, only the pixels in the mask are sampled to be predicted by the trained synapse segmentation model; otherwise, all pixels are subject to prediction. The local and probability features are extracted similarly to the training step, and pixels outside of the neurite mask are automatically classified as background.

#### Training data

One or more pairs of a fluorescent microscopy image and its corresponding binary label image are required to train the model (**Figure 2**). For the pre-trained models and strain-specific models used in this work, the pencil tool in Fiji (Schindelin et al., 2012) was used to generate binary label images.

#### Training of the first layer

The first-layer classifier is trained through two sequential training steps (**Figure 2 – figure supplement 2B**). In the first step, all positive pixels and a small portion of negative pixels near the positive ones are selected as a training set. Local features of these pixels are extracted and fed along with the label to train the first-layer classifier. Since only a small portion of negative pixels spatially close to the positive pixels is included in the initial training dataset, the first-layer classifier is particularly prone to type I error after this step. To resolve this issue, more negative pixels are added to the training dataset in the second step, and the classifier is retrained. After the first step, pixels in sampled patches are predicted by the trained classifier and false negative pixels are selected to be added to the training dataset. These target patches are sampled in a similar manner as before but are ten times larger in dimension. The first-layer classifier is then fitted again with this expanded training dataset.

#### Training of the second layer

As the size of a synapse is generally bigger than just a single pixel, a pixel adjacent to the positive foreground is more likely to be foreground than another pixel with the same local features but away from the foreground. To account for this factor, we added the probabilities of adjacent pixels being positive as additional features to the original local features for the second-layer classifier. For each pixel in the training dataset, probabilities of the target pixel and the six adjacent pixels (-1 and +1 in either x, y, or z direction) are calculated by the first-layer classifier and the feature vector is expanded with these six probability values. The second-layer classifier is then trained with the expanded features using the same model as the first-layer classifier (**Figure 2 – figure supplement 2B**).

#### Local features

The local features used in WormPsyQi’s synapse classifier include the pixel value, Sobel derivatives, difference of Gaussians (DoG), Laplacian of Gaussian (LoG), local average and standard deviation, and erosion of both synapse and neurite channels. Before training, each feature is normalized with a standard scaler, removing the mean and scaling to unit variance.

#### Patch sampling in training

Among the pixels in the label images, only a small portion of pixels in the patches surrounding positive labels is selected to be fed into the model as training data points (**Figure 2 – figure supplement 2B**). This step not only reduces the time and memory required for the training process but also results in higher accuracy by preventing the model from fitting to irrelevant data points. Each 2D slice in the original microscopy image is first divided into non-overlapping patches, and only the patches containing positive pixels in the corresponding patch in the label image are processed in the following steps. The size of the patches is set to twice the square root of the average area of the connected labels.

### Pre-trained synapse segmentation models

WormPsyQi provides four different pre-trained synapse segmentation models: 1) labeled as “GRASP (sparse)” in the GUI, trained with three *otIs612* (PHB>AVA GRASP) images, 2) “GRASP (dense)”, trained with three *otIs653* (ASK>AIA GRASP) images, 3) “CLA-1”, trained with six *otEx7503* (I5 CLA-1) images, and 4) “RAB-3”, trained with two *otEx7231* (ASK RAB-3). Random Forest model was used for all of them.

### Synapse distribution quantification

The relative locations of synaptic puncta along neurites were determined using synapse segmentation masks and neurite skeletons. The skeletons (**Figure 10**) were extracted on top of neurite segmentation masks generated by the pipeline, coupled with minor manual editing using NeuTube (Feng, Zhao, and Kim, 2015) in cases where the segmentation result was not perfect. For each synapse, its location was marked by the closest node of the skeleton with respect to its centroid coordinates. When a synapse was too large and unresolved, each pixel of the synapse was used instead of the centroid to give a smoother distribution. The distribution was normalized from 0 to 1 either by the entire neurite length (from soma to the axon terminal), or by synapse domain length (neurite segment occupied by synapses). The probability density of distribution of a population was acquired by combining all worms of the population and dividing the entire synaptic volume.

### Code availability

The WormPsyQi software is implemented with Python and is publicly available at https://github.com/lu-lab/worm-psyqi. PyTorch (Paszke et al., 2019) is used for implementing the deep CNN model in synapse segmentation process, and Scikit-learn (Pedregosa et al., 2011) package is mainly used for other methods. A comprehensive guide to installing and operating the software is on the Github repository. To facilitate use, the code is accompanied by a user manual with instructions for synapse quantification that is both standardized to yield quantitative data and flexible for customization.

## ACKNOWLEDGEMENTS

We thank Chi Chen for assistance with microinjections to generate strains; Ulkar Aghayeva for generating RAB-3 reporters for pharyngeal neurons MC, M3, and M4; Steven Cook, Emily Bayer, and Cyril Cros for kindly sharing previous image datasets for synapse analysis. This work was funded by an NSF NeuroNex award (#1707401) and NSF-Simons Southeast Center for Mathematics and Biology (#1764406).

## SUPPLEMENTAL TABLES

**Supplemental Table 1:** List of fluorescent reporters analyzed in this paper using WormPsyQi. Strain identifier, genotype, datasets analyzed, and imaging conditions (stage, sex) are listed for each reporter.

**Supplemental Table 2:** Details of transgenic constructs and injections.

**Supplemental Table 3:** Segmentation benchmark for different U-Net structures.

## EXTENDED FIGURE LEGENDS

**Figure 2 – figure supplement 1.**
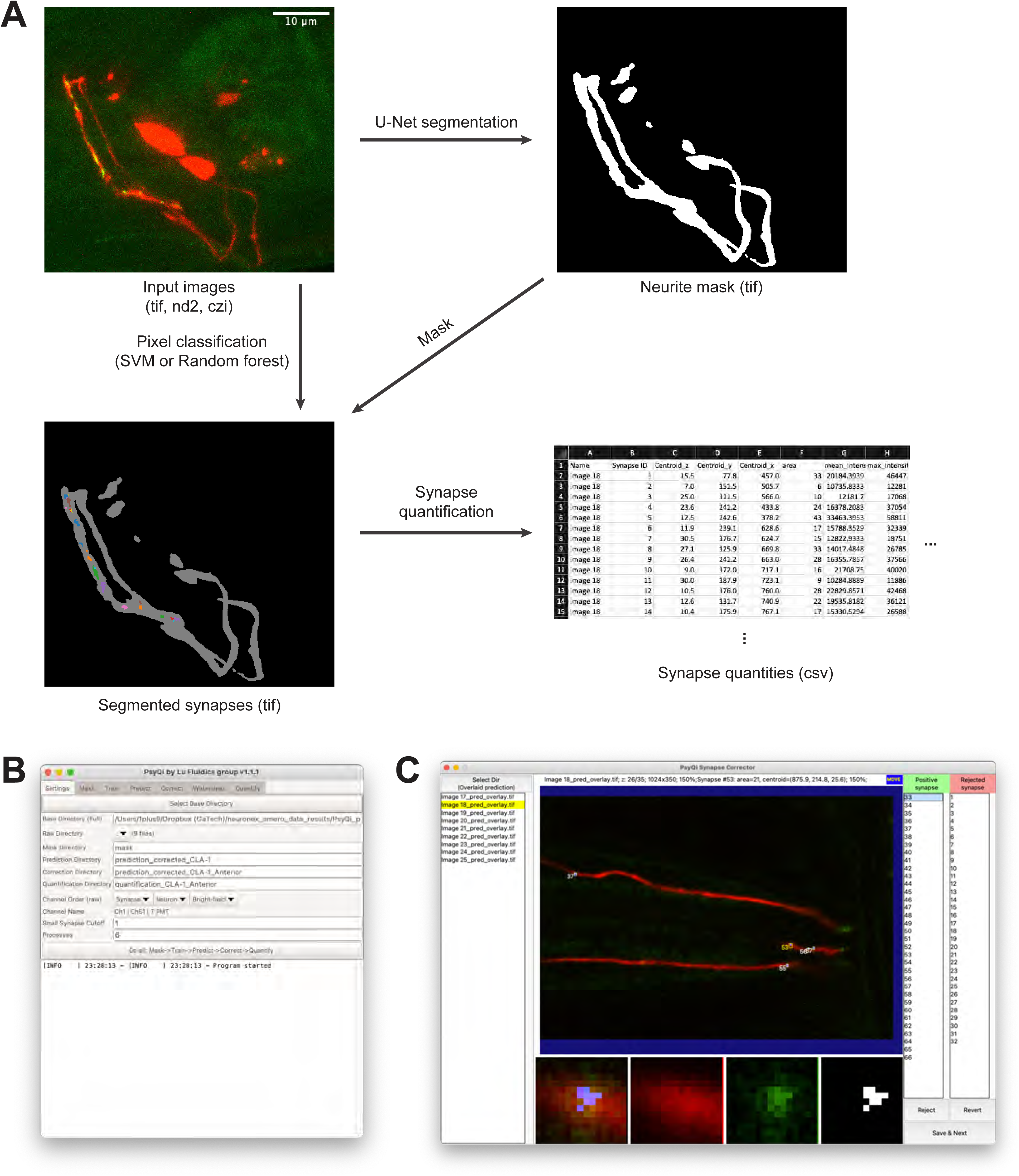
**(A)** WormPsyQi pipeline schematics with representative input and output. Input images are typically 3d multicolor stacks. Neurite segmentation process generates a 3d binary image labeling regions where synapses could be localized. Synapse segmentation process takes the input image and the neurite mask and generates a binary image labeling synapse. The example image is showing the connected components from the binary synapse label colored differently and overlaid with the neurite label in gray for visualization. Synapse quantification process extracts various synapse- and image-wise quantities and saves them as .csv format. **(B)** WormPsyQi main GUI. WormPsyQi provides a GUI to easily manage the processing parameters and the output data storage. **(C)** WormPsyQi GUI for reviewing and correcting synapse segmentation. Users can quickly scroll through the slices to visually inspect the quality of the synapse segmentation. False-positive synapses can be reviewed and rejected individually, or ROI can be selected to reject synapses in/out of the ROI.

**Figure 2 – figure supplement 2.**
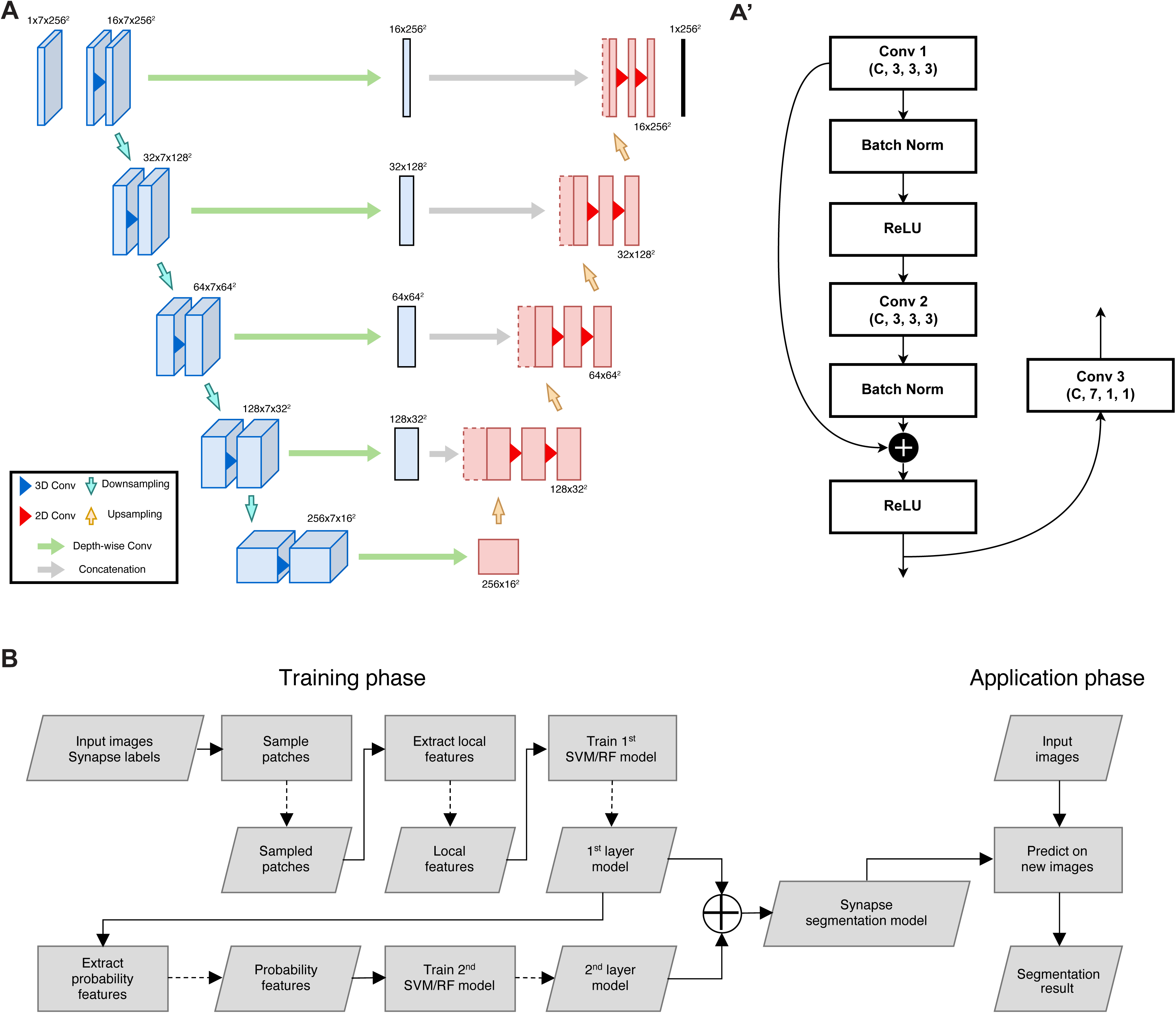
**(A)** 2.5D U-Net architecture for neurite segmentation. The input is a chunk with seven slices and the output is the segmentation mask of the center slice, with intermediate convolution blocks that combine depth-wise spatial context in the concatenation path. The structure of the encoding block is detailed in panel A’. **(B)** Flowchart for synapse segmentation. Rectangles represent a process or action, and parallelograms represent an input, output, or intermediate data.

**Figure 2 – figure supplement 3:**
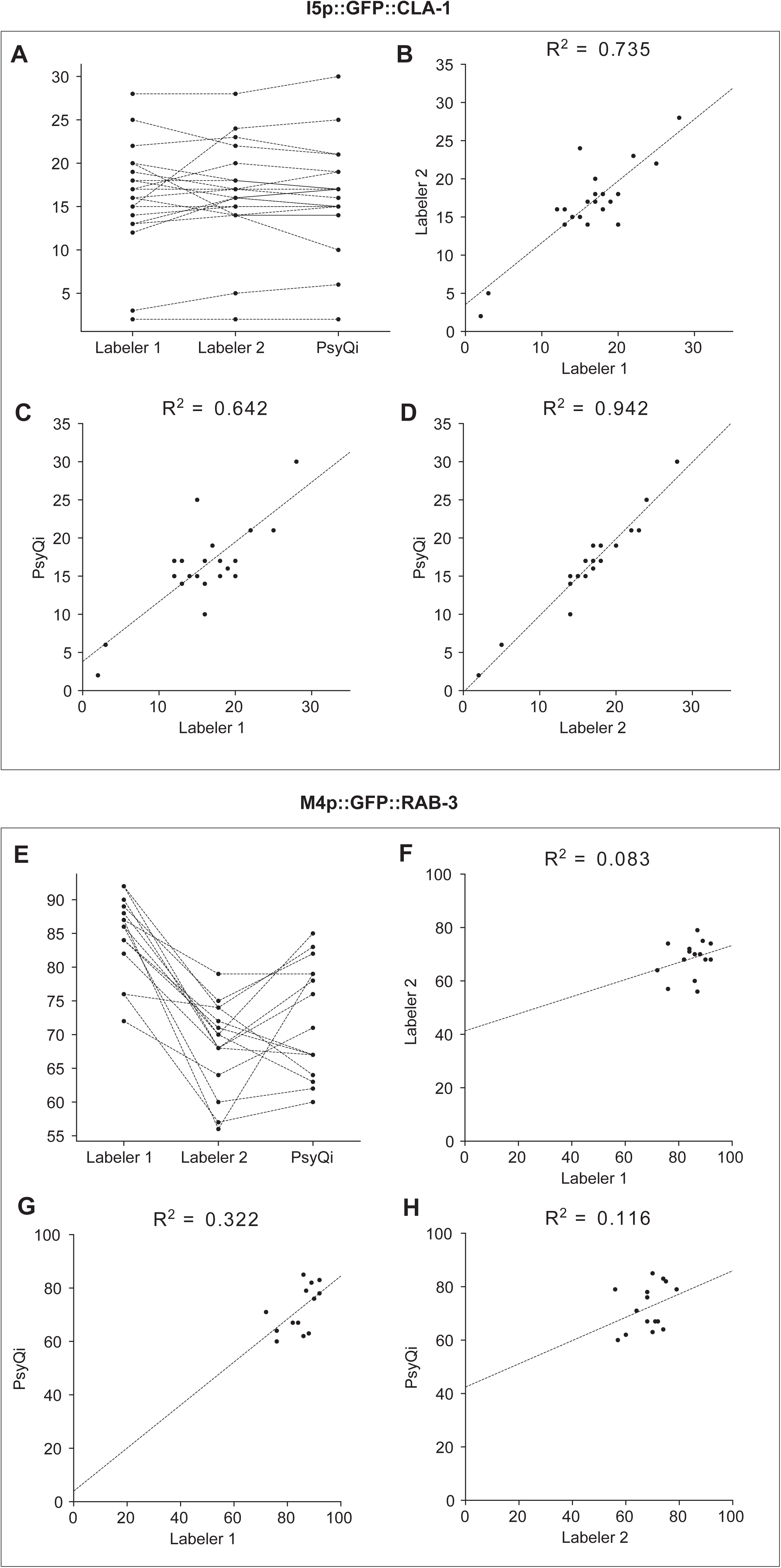
The correlation between WormPsyQi and manual count is comparable to the human-to-human correlation. Graphs depict the synapse count of individual worms for I5p::GFP::CLA-1 strain (*otEx7403*) **(A-D)** and M4p::GFP::RAB-3 strain (*otIs597*) **(E-H)**. WormPsyQi’s puncta count display a stronger or comparable correlation with manual counts than that seen between manual counts of two labelers, for both the good quality strain I5p::GFP::CLA-1 and the poor quality strain M4p::GFP::RAB-3. Each dot in the graphs represents an individual worm. **(A, E)** Changes in synapse count between two human labelers and WormPsyQi. Each dotted line is connecting counts from the same individual worm. **(B-D, F-H)** Pairwise synapse count comparison between **(B, F)** labeler 1 and labeler 2, **(C, G)** labeler 1 and WormPsyQi, and **(D, H)** labeler 2 and WormPsyQi. The dotted line represents the linear regression, and the Pearson correlation coefficient is displayed at the top of each graph.

**Figure 2 – figure supplement 4:**
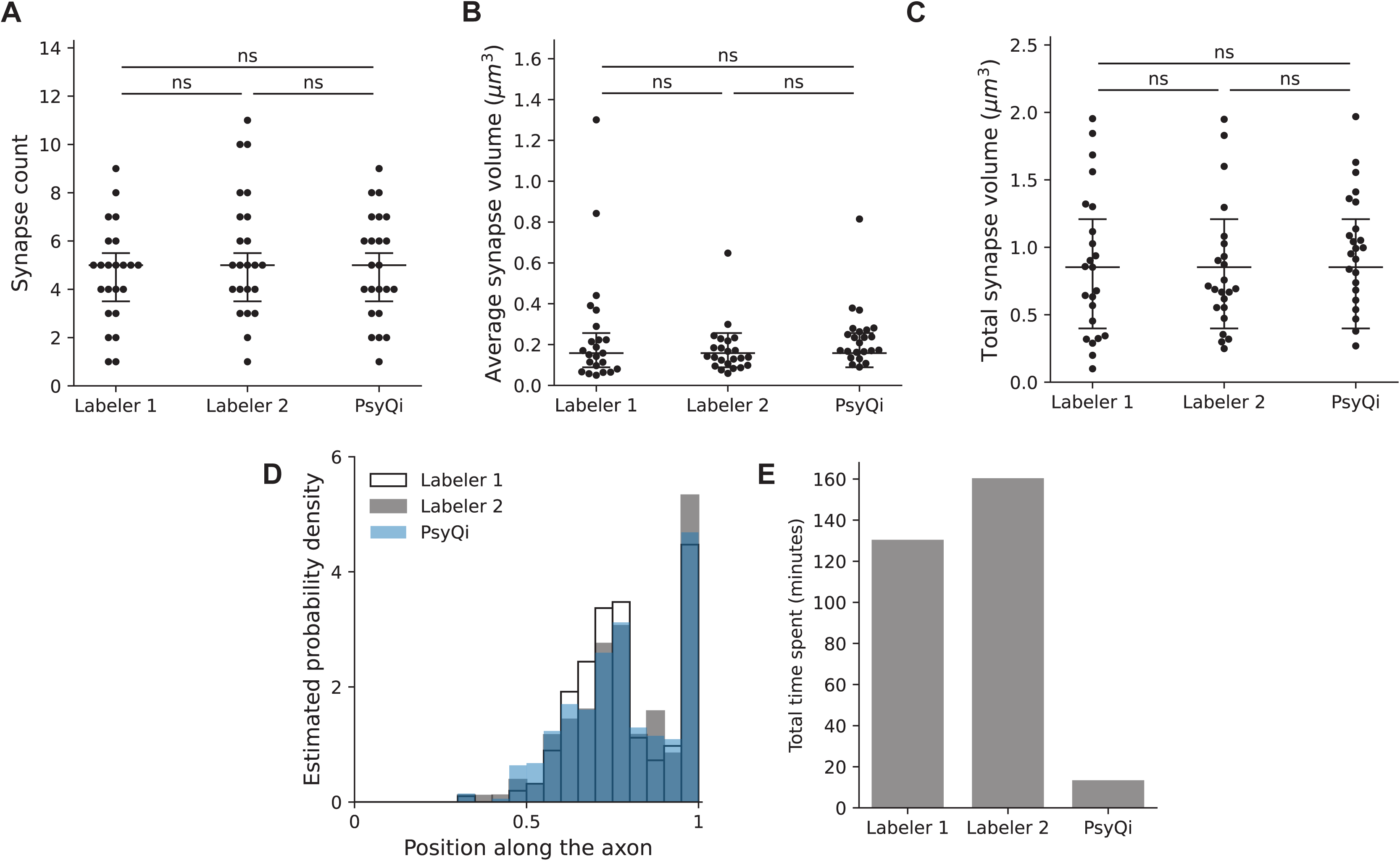
Comparison between WormPsyQi and human quantification for puncta features and processing time. For proof-of-principle, all quantification is based on an ASKp:::GFP::CLA-1 reporter (*otIs789*) **(A-C)** No statistically significant difference was observed in human vs. WormPsyQi scoring of total puncta (synapse) count **(A)**, average synapse volume **(B)**, and total synapse volume **(C). (D)** WormPsyQi’s segmentation results show a spatial distribution of synapses along the axon similar to the annotations of two individual human labelers. **(E)** WormPsyQi achieves over a 10-fold reduction in processing time: while Labelers 1 and 2 took 130 minutes and 160 minutes to annotate puncta, respectively, WormPsyQi processed the same dataset (15 images) in 13 minutes using an affordable desktop machine. Each dot on the graphs represents an individual worm.

**Figure 4 – figure supplement 1:**
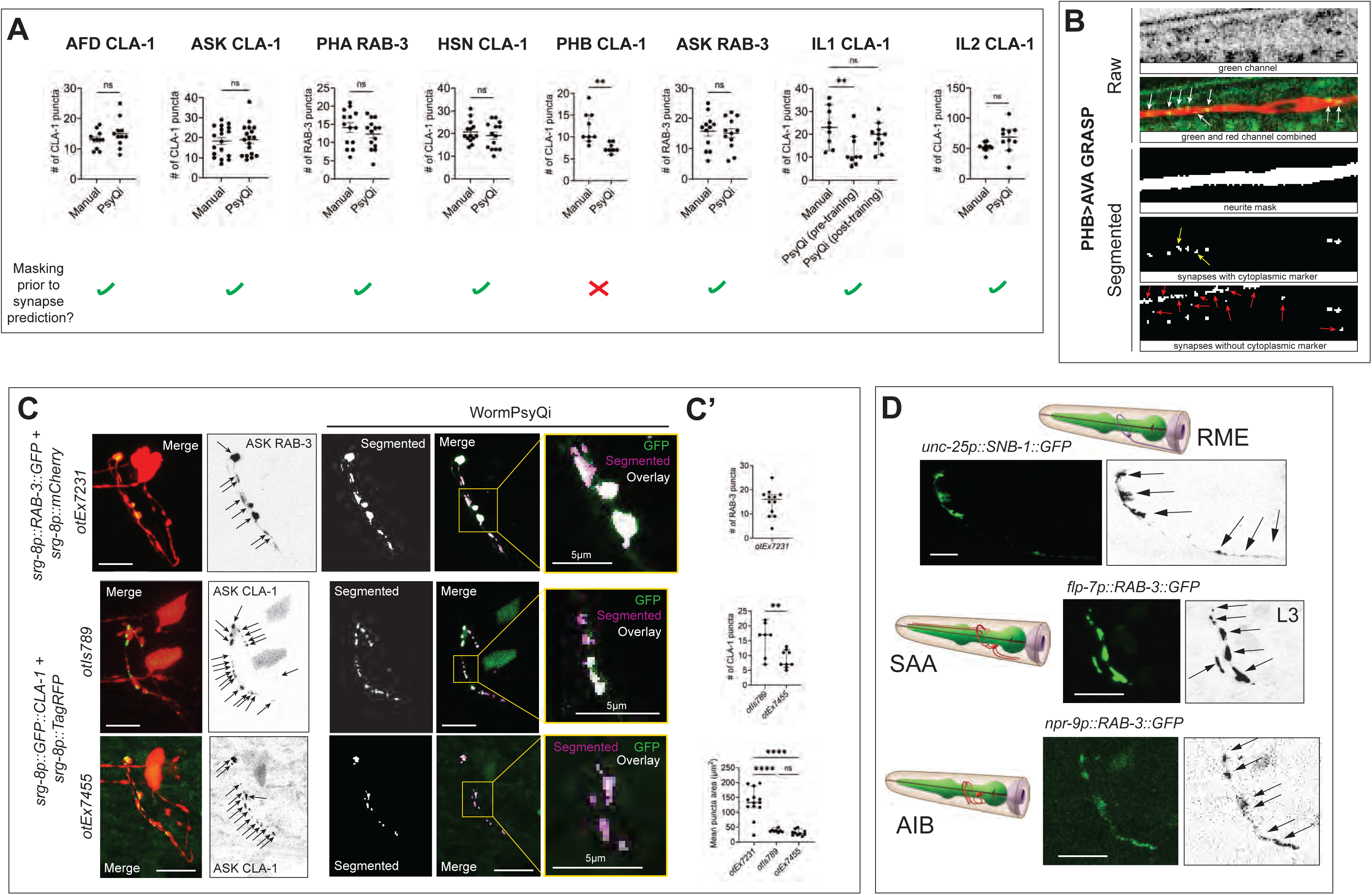
WormPsyQi performs optimally on images with a cytoplasmic marker and discrete puncta. **(A)** Manual vs. WormPsyQi quantification for representative GFP::CLA-1 (for neurons AFD, ASK, HSN, PHB, IL1 and IL2) and GFP::RAB-3 (for neurons PHA and ASK) reporters. No significant difference was observed in all cases besides PHB, where the masking step was skipped. The bottom row denotes whether a neurite mask was used prior to synapse segmentation in the pipeline. **(B)** Representative images of PHB>AVA GRASP synapse segmentation with and without the channel having cytoplasmic marker. The panels illustrate raw confocal images (top) and segmented neurite and synapses (bottom). The third panel displays WormPsyQi’s mask segmentation, the fourth panel shows synapse segmentation using the original image, and the last panel shows synapse segmentation using only the green channel (no-mask). The cytoplasmic marker not only helps exclude false-positive gut autofluorescence puncta (red arrows), but also assists in identifying some GRASP puncta that were missed in the no-mask segmentation (yellow arrows). All real GRASP puncta are pointed with white arrows. **(C)** Representative raw and segmented images and quantification of RAB-3 (*otEx7231*) CLA-1 reporters (*otIs789* and *otEx7455*) for ASK neurons. ASK presynaptic specializations were diffuse compared to both CLA-1 reporters (magnified insets), as denoted by mean puncta area, although relative number of puncta at L4 stage were similar **(C’)**. *P* values were calculated using an unpaired t-test or one-way ANOVA with Bonferroni correction. \*\*\*\**P* ≤ 0.0001, \*\**P* ≤ 0.01 and *P* > 0.05 not significant (ns). In each dataset in **(A)** and **(C)**, a dot represents a single worm and lines represent median with 95% confidence interval. **(D)** Images of a representative subset of reporters for neurons RME, SAA, and AIB, for which WormPsyQi could not segment and quantify puncta because of diffuse signal and high synapse density. Specializations were visualized using fluorescently-tagged presynaptic SNB-1 and RAB-3. All images are maximum intensity projections of confocal Z-stacks. Scale bars = 10μm unless otherwise noted.

**Figure 6 – figure supplement 1:**
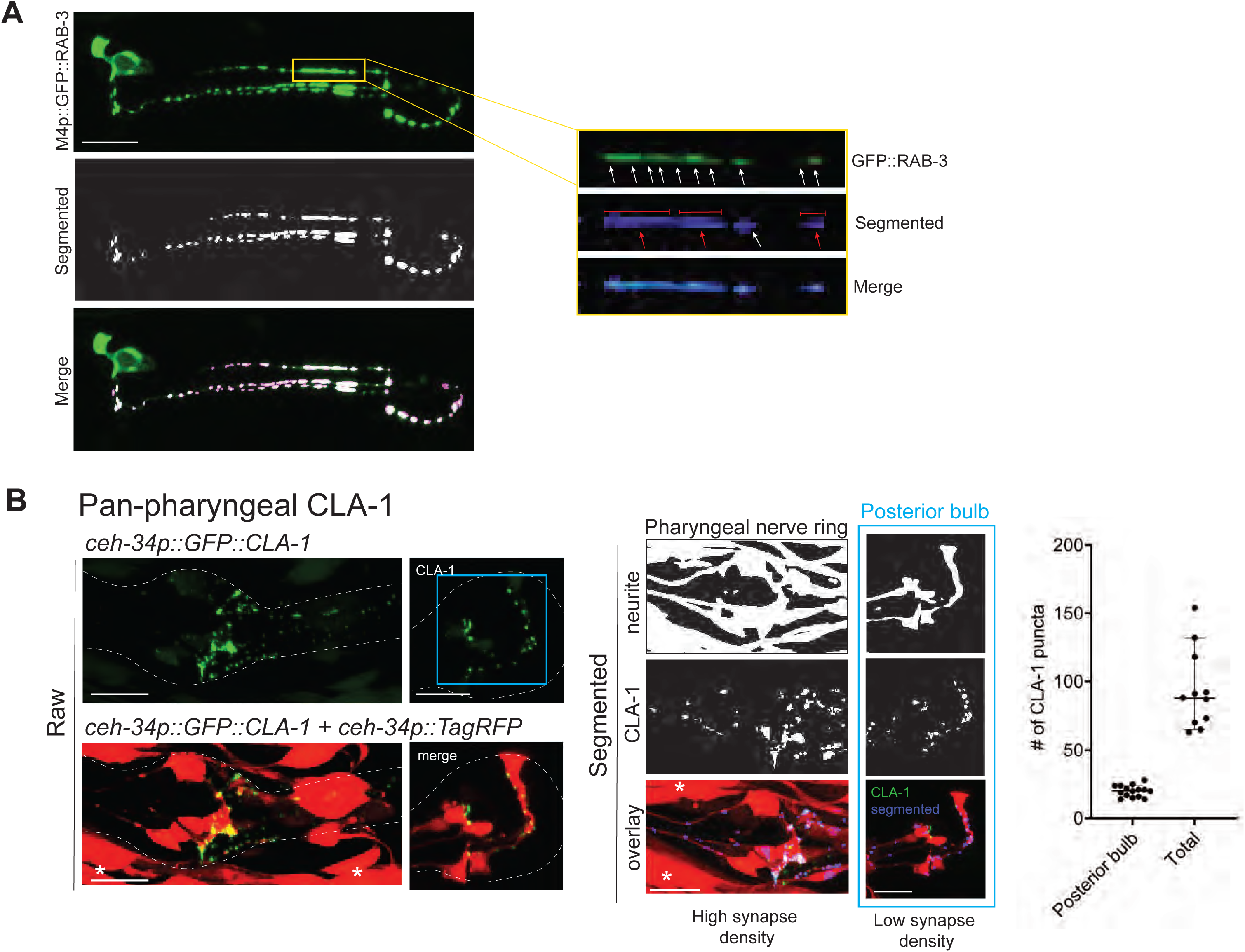
WormPsyQi analysis of all presynaptic specializations in the pharynx. **(A)** Representative images of M4p::GFP::RAB-3 (*otIs597*) processed with WormPsyQi. Magnified insets show a synapse-dense region where multiple puncta (white arrows) are discernible in the raw GFP image, but WormPsyQi segments adjacent puncta together (red arrows), therefore undercounting as presented in the larger dataset for the same reporter in Figure 6F. **(B)** Representative images and quantification of a pan-pharyngeal synaptic reporter (*otIs785*). Neuronal processes were labeled with *ceh- 34p::TagRFP* and presynaptic specializations were visualized with *ceh-34p::GFP::CLA-*1 Raw (top), segmented (middle), and overlap (bottom) images are shown. Quantification of puncta in the whole pharynx (‘Total’) and posterior pharyngeal bulb (blue) are shown in the graph. In each dataset, a dot represents a single worm and lines represent median. Compared to the posterior bulb the spread is greater for total count, where synapse segmentation was suboptimal due to a high synapse density and inseparable processes. All images are maximum intensity projections of confocal Z-stacks. Muscle expression from *ceh-34p* is marked with an asterisk. Scale bars = 10μm.

**Figure 9 – figure supplement 1:**
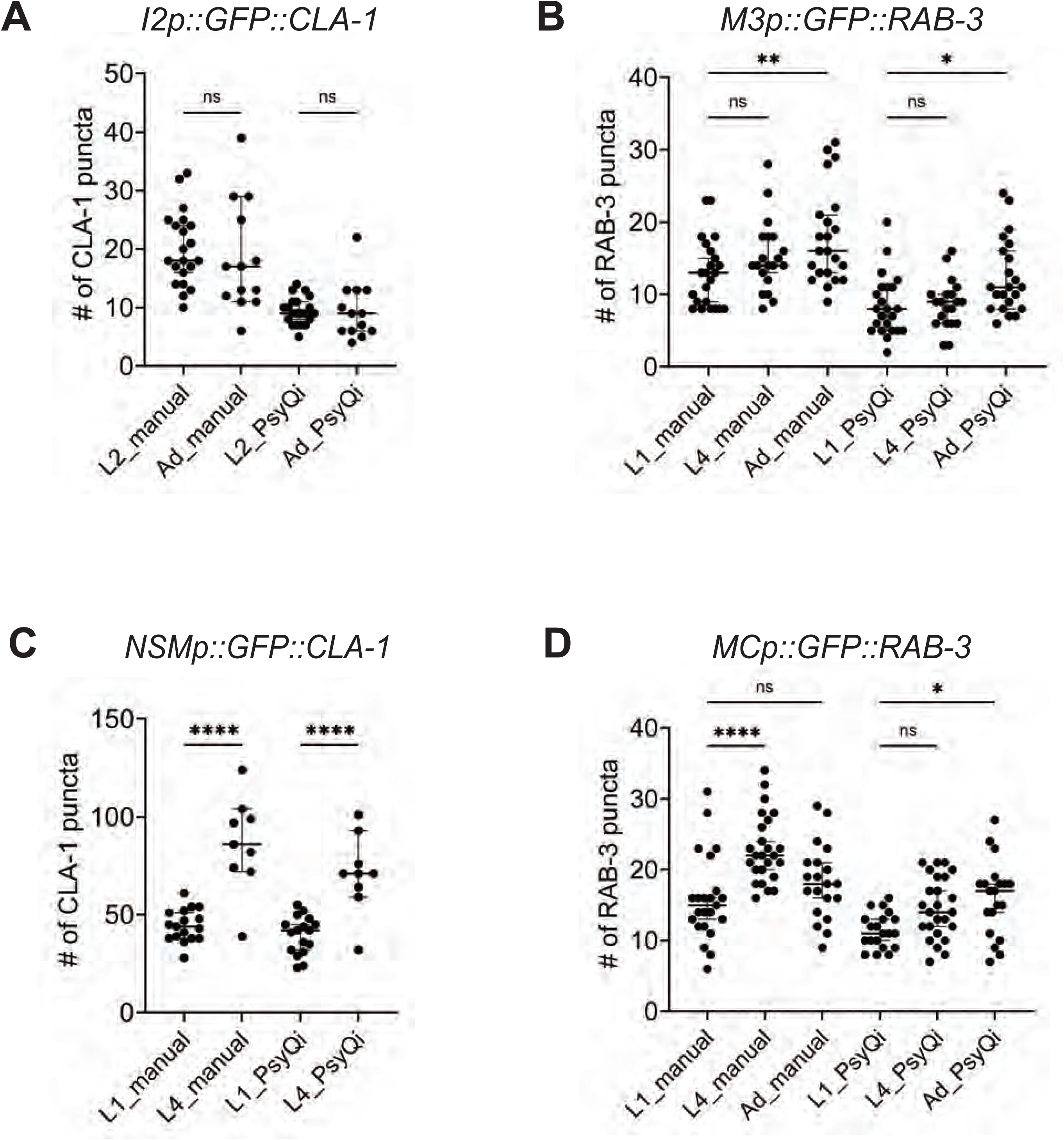
Manual and WormPsyQi quantification show the same developmental trends in synapses in the pharynx. Graphs showing manual vs. WormPsyQi puncta quantification for presynaptic specializations in pharyngeal neurons **(A)** I2, **(B)** M3, **(C)** NSM, and **(D)** MC in early larval (L1, L2) and late larval (L4) or adult (Ad) stages. Although WormPsyQi scores fewer puncta relative to manual counts in L1 and L2 animals, the trends between the two modes of scoring are consistent with modest synapse addition in neurons I2, M3, and MC, and significant addition in neurons NSM. *P* values were calculated using one-way ANOVA with Bonferroni correction. \*\*\*\**P* ≤ 0.0001, \*\**P* ≤ 0.01, \**P* ≤ 0.05 and *P* > 0.05 not significant (ns). In each dataset, a dot represents a single worm and lines represent median with 95% confidence interval.

**Figure 9 – figure supplement 2:**
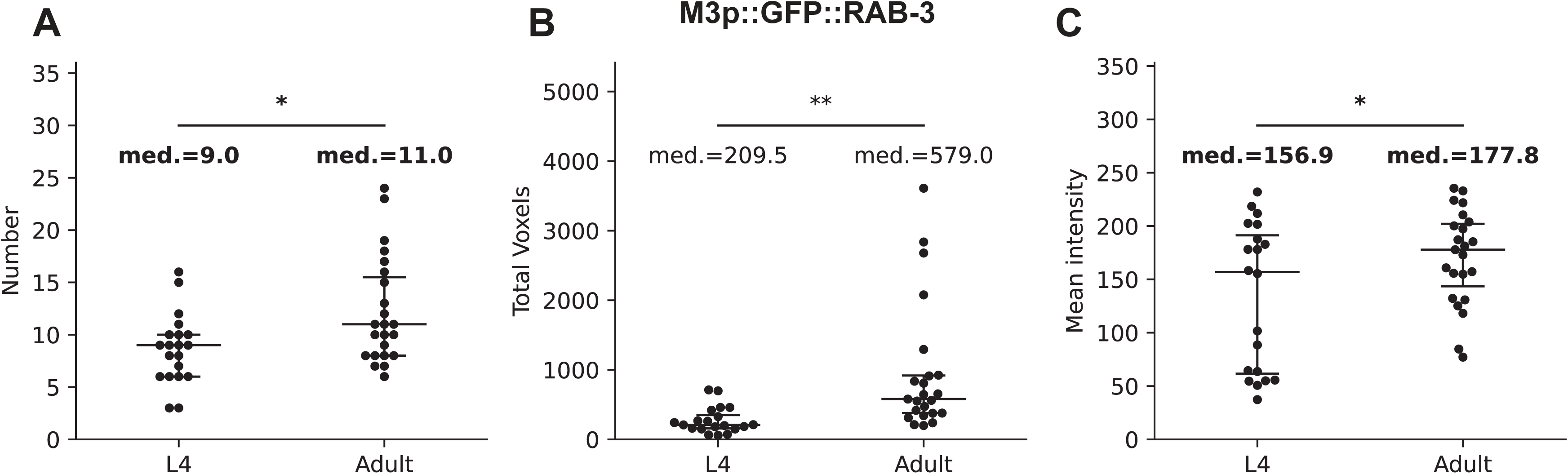
WormPsyQi quantification reveals small but significant differences between adjacent developmental stages. Graphs showing the comparison of WormPsyQi quantification on synapse count **(A)**, total puncta volume **(B)**, and mean pixel intensity of puncta **(C)** for M3p::GFP::RAB-3 reporter (*otIs602*) between L4 and adult stages. All three quantities show significant changes from L4 to adult, even when the changes are marginal (10∼20%) in the case of total puncta count and mean pixel intensity (highlighted in bold). *P* values were calculated using one-way ANOVA.

**Figure 10 – figure supplement 1:**
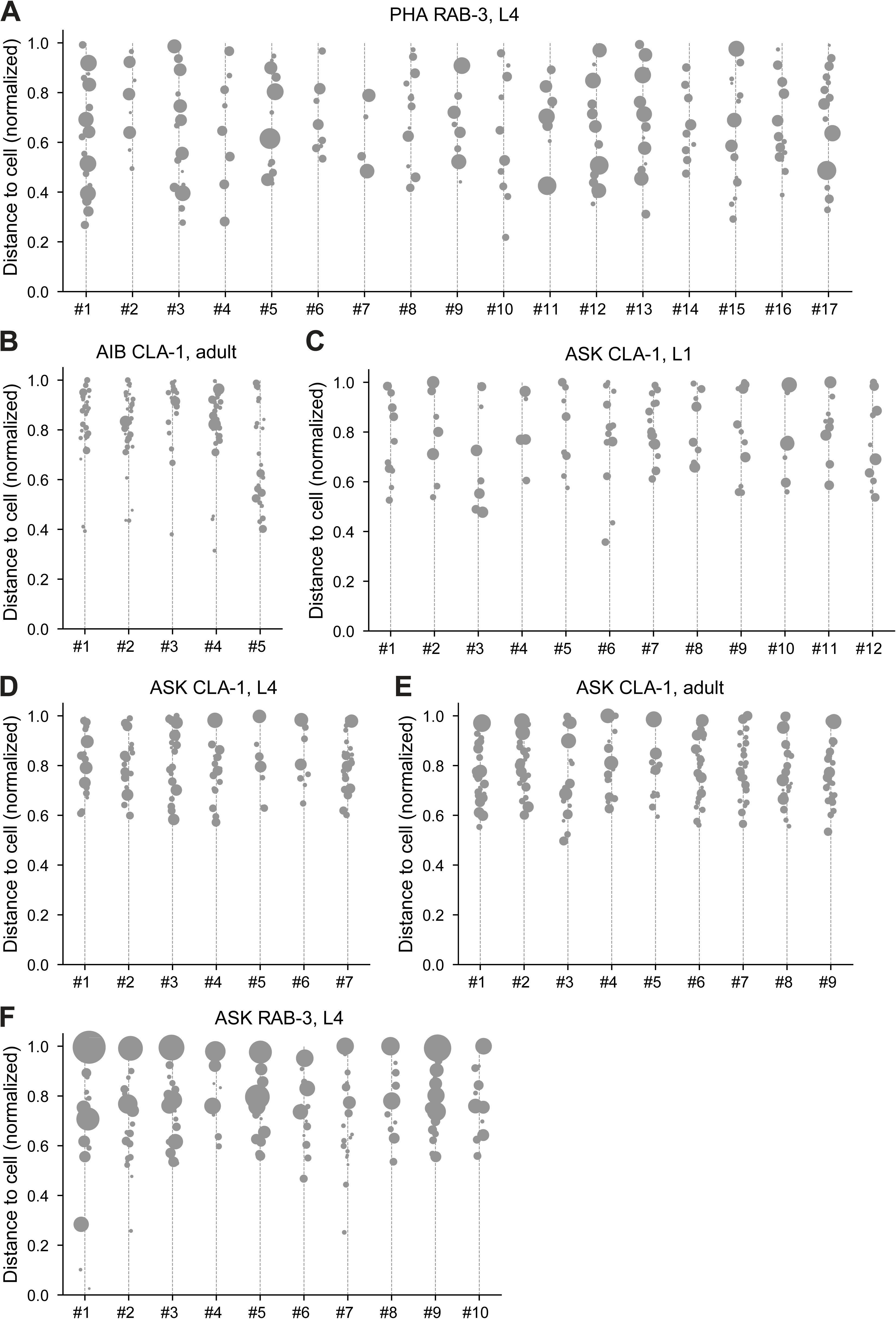
Distribution of CLA-1 puncta in individual worms. Individual CLA-1 puncta distribution in cell-specific reporters for neurons PHA, AIB, and ASK (population-level quantification and statistics summarized in Figure 10). Each column in the graphs corresponds to an individual worm. Gray dots denote discrete puncta, and the size of dots corresponds to the synapse volume. The y-axis represents the normalized distance from the soma, with 0 being the closest and 1 being the farthest point in the region of interest. **(A)** PHA RAB-3 adults, **(B)** AIB CLA-1 L4s, **(C)** ASK CLA- 1 L1s, **(D)** ASK CLA-1 L4s, **(E)** ASK RAB-3 L4s, and **(F)** ASK CLA-1 adults.

**Figure 10 – figure supplement 2:**
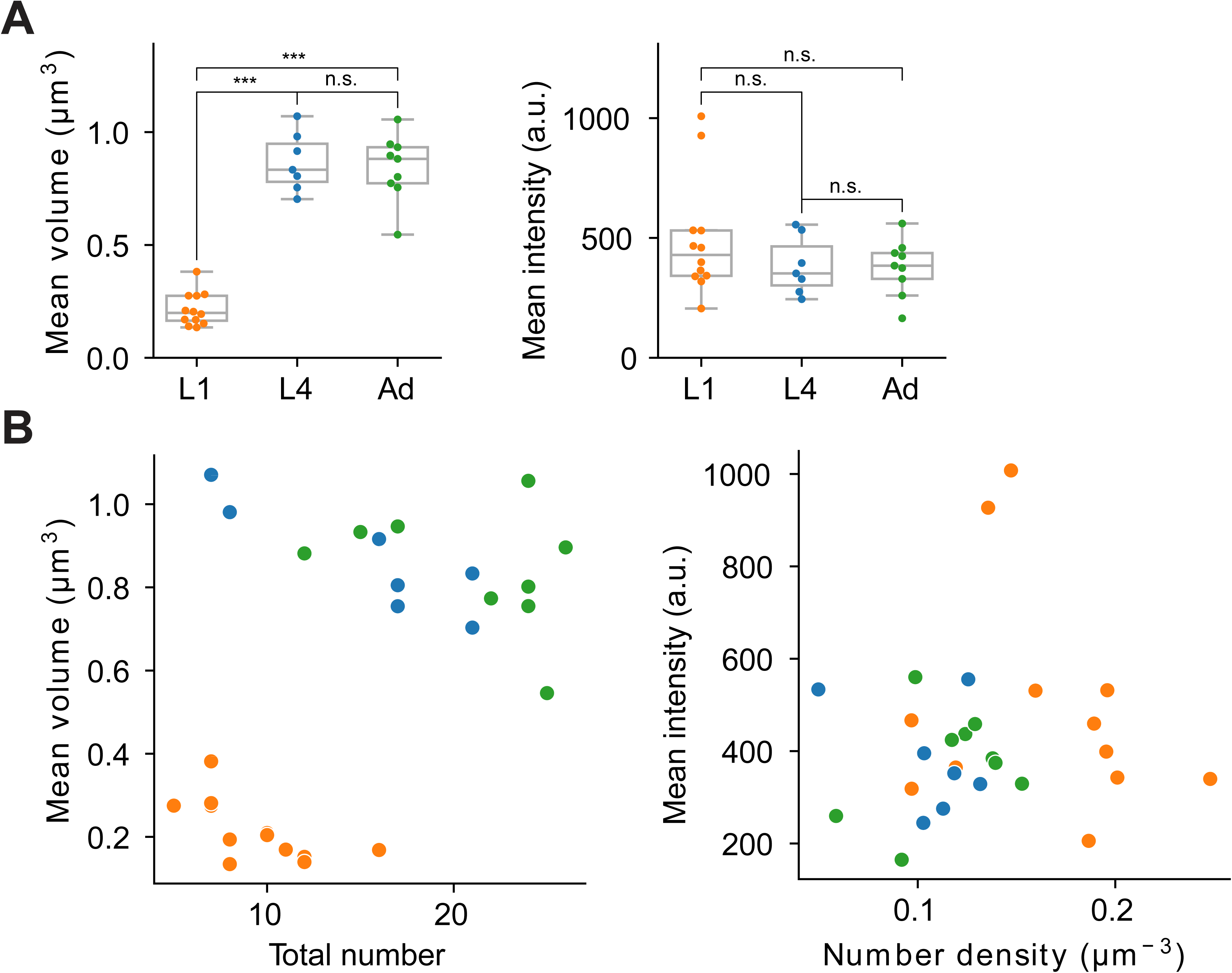
Extended features and representative two-dimensional scatter plots of ASK CLA-1 reporter across different developmental stages. **(A)** Left: Average volume of synaptic puncta. Right: Average fluorescence intensity. Each dot in a dataset represents a single worm. *P* values were calculated using one-way ANOVA with Bonferroni correction for multiple comparisons \*\*\**P* ≤ 0.001, and *P* > 0.05 not significant (ns). **(B)** Two examples illustrating two-dimensional synapse ultrastructure analysis. Color coding is consistent with Figure 10. Left: Total number vs. average volume of puncta at L1, L4, and adult stages. In this representation, L1 stage worms are clustered separately from L4 and adult stages, which are grouped together. Right: Number density vs. average intensity of puncta; developmental stages are not clearly differentiated in this feature space.

